# Effect of Climate Warming on Mosquito Population Dynamics in Newfoundland

**DOI:** 10.1101/2025.09.07.674692

**Authors:** Joseph Baafi, Amy Hurford

**Author notes:** Corresponding author’s. Contributing author’s. These authors contributed equally to this work.

## Abstract

Mosquitoes are key vectors of several infectious diseases affecting humans and animals. In North America, *Culex* mosquitoes are primary vectors of West Nile virus, St. Louis encephalitis, and Japanese encephalitis, as well as pathogens that impact birds and horses. The *Culex* life cycle consists of four stages, eggs, larvae, pupae, and adults, each with distinct development and mortality rates. Only active (non-diapausing) adults can reproduce, and environmental factors such as temperature, photoperiod, and rainfall influence population dynamics and stage-specific abundances. We develop a stage-structured model that integrates historical climate data and available experimental data to describe how key climate variables regulate life history parameters. Specifically, oviposition rates depend on temperature, maturation and survival are influenced by both temperature and rainfall, mortality is modeled as temperature-dependent, and diapause induction and termination are driven by photoperiod. Unlike many previous models focused on tropical mosquitoes, our framework explicitly incorporates diapause, a critical adaptation for temperate *Culex* populations. Simulations reveal strong nonlinear responses to warming, moderate temperature shifts amplify the differences in mosquito abundance across rainfall regimes, whereas higher warming leads to convergence at consistently high densities. Rainfall amounts determine whether populations remain suppressed, variable, or strongly amplified, whereas warming extends the active season and elevates population abundance. These results underscore the importance of jointly considering temperature, rainfall, and photoperiod in predicting mosquito dynamics. Our findings suggest that climate change may expand the seasonal window of mosquito activity and raise vector abundance, thereby increasing opportunities for pathogen transmission. The model provides a mechanistic framework for exploring how interacting climate drivers shape vector ecology and highlights priorities for data collection and adaptive control strategies under environmental change.

## Introduction

Mosquitoes are major vectors of infectious diseases affecting millions of people worldwide (World Health Organization, 2025; World Health Organization, 2023; Gubler, 2011; Weaver et al., 2018; Tolle, 2009; Bhatt et al., 2013). In 2024, the Americas experienced their worst recorded dengue outbreak, with over 13 million cases reported across North, Central, and South America and the Caribbean (Pan American Health Organization, 2024; World Health Organization, 2025; Pan American Health Organization, 2025). In the United States, West Nile virus (WNV) remains the most prevalent mosquito-borne disease, with 1,791 reported cases and 164 deaths in 2024, more than 7% of which were neuroinvasive (Centers for Disease Control and Prevention, 2025). The Centers for Disease Control and Prevention (CDC) estimates that approximately 20% of WNV-infected individuals develop symptoms, while fewer than 1% experience severe neuroinvasive disease such as meningitis or encephalitis (Centers for Disease Control and Prevention, 2025; Petersen, Brault, and Nasci, 2013). Although Eastern Equine Encephalitis (EEE) cases remain rare (19 reported in 2024), its case fatality rate averages around 30%(Centers for Disease Control and Prevention, 2024; Lindsey, Staples, and Fischer, 2018). In temperate regions of North America, *Culex* mosquitoes, particularly *Culex pipiens* and *Culex quinquefasciatus* are the primary vectors of WNV(Centers for Disease Control and Prevention, 2025; Farajollahi et al., 2011), while *Aedes* species drive the transmission of dengue, Zika, and other arboviruses in many tropical and subtropical regions (World Health Organization, 2025; Pan American Health Organization, 2024). With no specific treatments available for most of these diseases (Pan American Health Organization, 2024), controlling vector populations remains the cornerstone of prevention in endemic areas (Pan American Health Organization, 2025; World Health Organization, 2025). The abundance and distribution of these vectors are shaped by complex interactions between biological traits and environmental conditions, with weather and climate playing a central role (Joseph Baafi and Hurford, 2025; Beck-Johnson et al., 2013; Abdelrazec and Gumel, 2017; Frantz et al., 2024; Ewing et al., 2016).

Weather variables such as temperature (Reinhold, Lazzari, and Lahondère, 2018; Winokur et al., 2020), rainfall (Beck-Johnson et al., 2013), and photoperiod (Diniz et al., 2017) strongly influence mosquito life-history traits, including development (Ciota, Matacchiero, et al., 2014), reproduction (Coon, Hegde, and Hughes, 2022), and survival (Winokur et al., 2020). Warmer temperatures often accelerate development and extend transmission seasons (Coon, Hegde, and Hughes, 2022; Reisen, 2013), while rainfall governs the availability and stability of breeding habitats, both drought and excessive precipitation can limit population growth and alter seasonal timing (Beck-Johnson et al., 2013). Photoperiod regulates reproductive cycles, with shorter days inducing diapause in some *Culex* species, enabling persistence through unfavorable conditions (Diniz et al., 2017). The combined and often nonlinear effects of these drivers shape mosquito phenology and abundance (Reinhold, Lazzari, and Lahondère, 2018; Ciota and Keyel, 2019), with elevated temperatures potentially boosting vector densities, but extreme events creating survival bottlenecks (Abdelrazec and Gumel, 2017). Identifying the thresholds at which these factors affect life stages and abundance is critical for predicting mosquito responses to climate warming (Reinhold, Lazzari, and Lahondère, 2018; Beck-Johnson et al., 2013; Joseph Baafi and Hurford, 2025). Such complexity underscores the need for mathematical modeling approaches capable of integrating multiple environmental drivers and capturing their interactions.

A wide range of mathematical models have been developed to investigate mosquito population dynamics under varying environmental conditions (Ezanno et al., 2015; Beck-Johnson et al., 2013; Joseph Baafi and Hurford, 2025; Ewing et al., 2016). Recent approaches include weather-driven ordinary differential equation (ODE) models that integrate temperature, photoperiod, intra-species competition, and habitat variability to simulate species such as *Culex pipiens* (Bhowmick et al., 2025; Frantz et al., 2024; Abdelrazec and Gumel, 2017). Other studies have used partial differential equation (PDE) models incorporating seasonal environmental variation to capture the dynamics of different mosquito species(Beck-Johnson et al., 2013; Ewing et al., 2016). These frameworks support the forecasting of vector abundance and seasonal activity to guide vector control strategies (Beck-Johnson et al., 2013). Some models have incorporated overwintering mechanisms (Ewing et al., 2016; Frantz et al., 2024), recognizing that behaviors such as diapause are essential for capturing population persistence and the seasonal carryover of disease risk, particularly in temperate regions (Diniz et al., 2017). While these models have advanced our ability to predict abundance, gaps remain in representing ecological mechanisms like diapause and in quantifying how ongoing climate change will influence mosquito dynamics and associated disease risks (Reinhold, Lazzari, and Lahondère, 2018).

We develop a mechanistic, stage-structured model for *Culex* mosquito population dynamics that explicitly incorporates adult diapause, an ecological mechanism rarely parameterized in climate-driven mosquito models. Environmental forcing couples empirically informed responses to three key drivers, temperature, which modulates development, fecundity, and survival (Bhowmick et al., 2025; Reinhold, Lazzari, and Lahondère, 2018; Mordecai et al., 2019); photoperiod, which governs diapause entry and termination (Diniz et al., 2017); and rainfall, which regulates aquatic habitat availability and early-stage survival (Beck-Johnson et al., 2013; Reiter, 2001). To capture hydroclimatic variability, rainfall is simulated as a stochastic process while preserving its seasonal pattern. Using observed seasonal climate, along with prescribed warming and modified-rainfall scenarios, we quantify changes in abundance and phenology, including the onset and duration of the active season and peak abundance. By integrating overwintering dynamics with multiple seasonal drivers in a unified framework, our approach captures nonlinear responses and ecological thresholds that are often overlooked. This novelty enables improved identification of climate-sensitive periods for targeted vector control and public health planning, particularly under future climate change scenarios.

The remainder of the paper is organized as follows: Section 2 describes the stage-structured mosquito modeling framework, data sources, climate-scenario design and computational methods. Section 3 presents the simulation results, and Section 4 discusses their implications for vector control and public health. The conclusion of the study is presented in Section 5.

## 2 Methods

### 2.1 Model structure

The mosquito life cycle consists of four main stages: egg, larva, pupa, and adult. In *Culex* mosquitoes, adult females deposit rafts of eggs on the surface of standing water. These eggs hatch into larvae, which pass through several instars before transforming into pupae. Pupae then undergo metamorphosis to emerge as adults, females seek vertebrate hosts for blood meals, enabling egg production and continuation of the cycle (Ewing et al., 2016; Joseph Baafi and Hurford, 2025).

In many temperate species, including *Culex* spp., population persistence through winter is achieved via diapause in inseminated adult females (Ewing et al., 2016; Frantz et al., 2024). This dormant state is typically triggered by seasonal cues such as photoperiod and temperature, allowing survival during periods unsuitable for reproduction and development.

We represent the life cycle using a system of stage-structured ordinary differential equations (ODEs) that track the abundances of eggs *E*(*t*), larvae *L*(*t*), pupae *P* (*t*), active adults *A*(*t*), and diapausing adults *A*_*d*_(*t*) at time *t*. The stage-structured ODE framework builds on the general life-cycle formulation described in Joseph Baafi and Hurford (2025), with additional structure to account for adult diapause following the approach of Frantz et al. (2024). While Frantz et al. (2024) implemented their model using partial differential equations, our formulation employs ODEs and introduces a key extension, all biological rates including development, survival, and reproduction are explicitly coupled to environmental drivers such as temperature, photoperiod, and rainfall. This extension enables the model to capture a wider range of environmentally induced variability than in previous formulations. An outline of the model is presented below, with full derivations provided in Appendix A.1. The state equations for *E*(*t*), *L*(*t*), *P* (*t*), *A*(*t*), and *A*_*d*_(*t*) are,

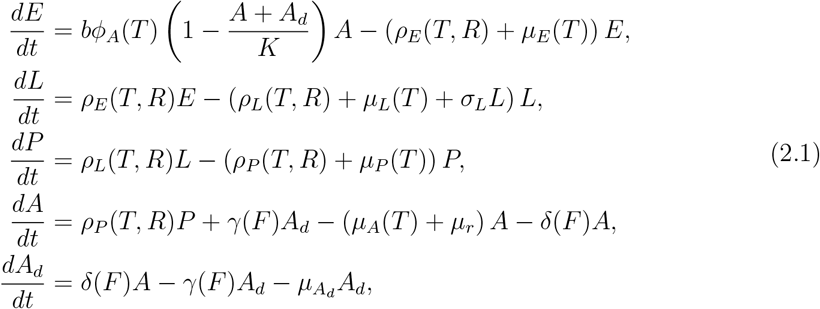

with initial conditions *E*(0), *L*(0), *P* (0), *A*(0), and *A*_*d*_(0). The model is driven by temperature *T* (*t*), rainfall *R*(*t*), and photoperiod *F* (*t*) at time *t*. We assume that immature and adult mosquitoes experience the same thermal environment, such that ambient air temperature approximates surface-water temperature (Joseph Baafi and Hurford, 2025; Abdelrazec and Gumel, 2017; Beck-Johnson et al., 2017; Beck-Johnson et al., 2013). The climate inputs *T* (*t*), *R*(*t*) and *F* (*t*)are taken as non-negative, continuous, bounded, and periodic functions. Stage-specific rates, *ϕ*_*A*_(*T*), *ρ*_*E*_(*T, R*), *ρ*_*L*_(*T, R*), *ρ*_*P*_ (*T, R*), *δ*(*F*), *γ*(*F*), *µ*_*E*_(*T*), *µ*_*L*_(*T*), *µ*_*P*_ (*T*), and *µ*_*A*_(*T*) are assumed to be non-negative, continuous, and bounded, with periodicity in time arising from the periodic environmental drivers.

In system (A.1), each term represents either the inflow to a stage via oviposition, hatching, or maturation, or the outflow via maturation to the next stage or mortality. The larval stage includes a density-dependent mortality term to capture intra- and inter-specific competition for food and habitat space, a feature commonly incorporated in mosquito population models (Joseph Baafi and Hurford, 2025; Lutambi et al., 2013; Okuneye, Abdelrazec, and Gumel, 2018). A schematic representation of the life-cycle transitions is shown in Figure 1 (Baafi, 2025).

**Figure 1.**
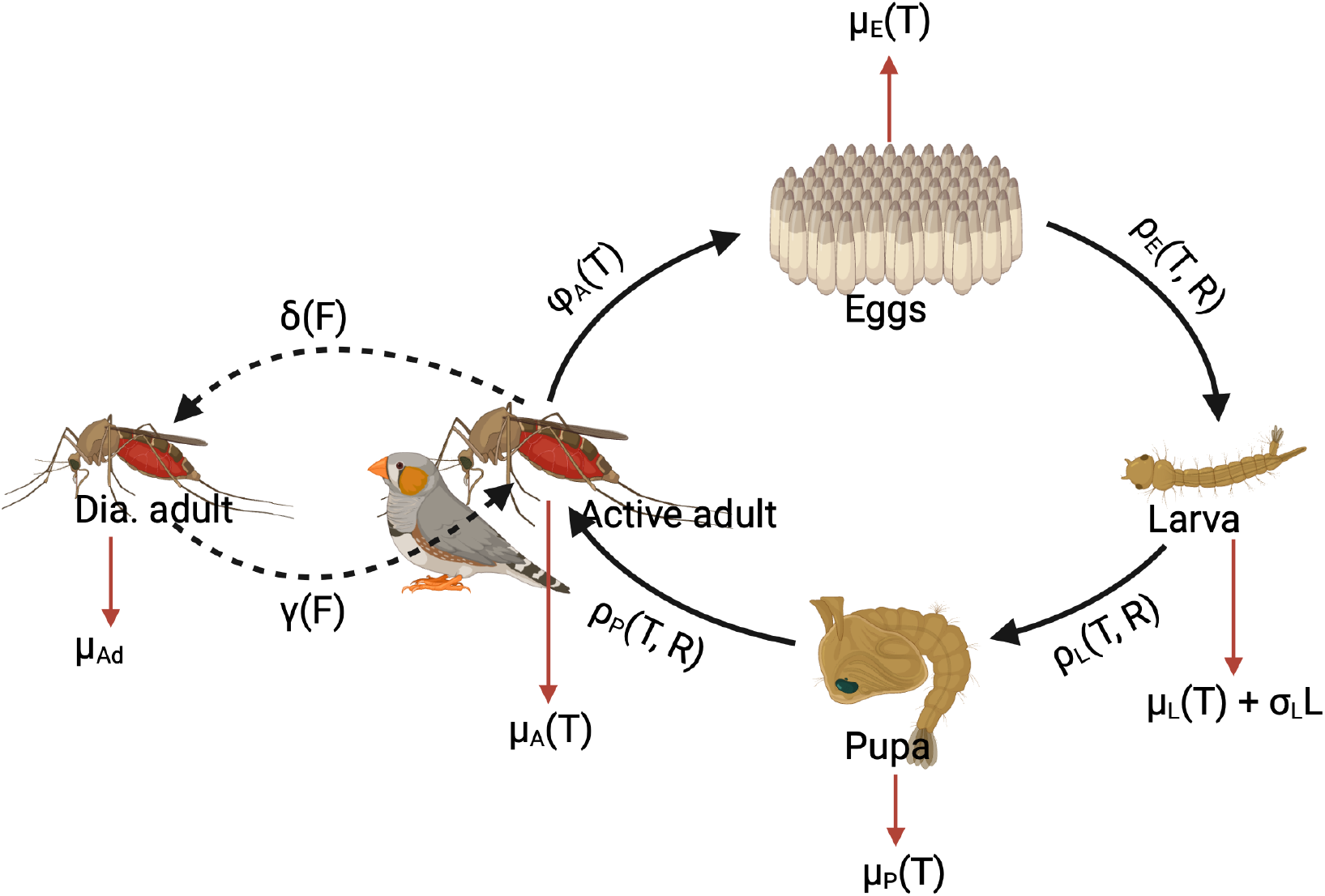
Schematic representation of the stage-structured *Culex* mosquito population model, created with BioRender.com (Baafi, 2025). The stage-structure consists of eggs (*E*), larvae (*L*), pupae (*P*), active adults (*A*), and diapausing adults (*A*_*d*_). Solid arrows represent oviposition (*ϕ*_*A*_) and stage transitions driven by development rates *ρ*_*E*_, *ρ*_*L*_, and *ρ*_*P*_ ; dashed arrows represent diapause entry (*δ*) and termination (*γ*). Environmental drivers (temperature (*T*), rainfall (*R*), and photoperiod (*F*)) influence stage-specific development, survival, reproduction, and diapause activation and termination rates.

The model is applied to examine *Culex* stage structure dynamics and to project changes in total abundance, active season duration, and peak abundance under climate-warming

and altered-rainfall scenarios. In contrast to earlier formulations, our framework explicitly incorporates adult diapause into a mechanistic stage-structured model, with modifications to stage-specific processes that build on but extend the formulation of Abdelrazec and Gumel (2017). By integrating temperature, photoperiod, and rainfall effects on multiple life-history rates, the model advances beyond previous work such as Jia et al. (2016), which considered only temperature and photoperiod. Furthermore, all biological rates are dynamically coupled to environmental drivers, providing an extension to Frantz et al. (2024), which modeled diapause but did not account for weather-driven variation across all rates. Collectively, these improvements enable a more realistic representation of seasonal and interannual variability in temperate mosquito populations. A complete list of model variables and parameters, including their units, sources, and environmental dependencies, is provided in Tables 1 and 2.

**Table 1.** Model variables (MVs), parameters (Params), definitions, and units.

| MVs/Params | Description | Unit |
| --- | --- | --- |
| $E(t)$ | Number of eggs at time $t$ | Number |
| $L(t)$ | Number of larvae at time $t$ | Number |
| $P(t)$ | Number of pupae at time $t$ | Number |
| $A(t)$ | Number of active adults at time $t$ | Number |
| $A_d(t)$ | Number of diapausing adults at time $t$ | Number |
| $b$ | Number of eggs laid per oviposition | Number |
| $\phi_A(T)$ | Eggs oviposition rate | female<sup>-1</sup> day<sup>-1</sup> |
| $K$ | Adult carrying capacity | Number |
| $\rho_E(T, R)$ | Egg development rate | day<sup>-1</sup> |
| $\rho_L(T, R)$ | Larval development rate | day<sup>-1</sup> |
| $\rho_P(T, R)$ | Pupal development rate | day<sup>-1</sup> |
| $\mu_E(T)$ | Egg mortality rate | day<sup>-1</sup> |
| $\mu_L(T)$ | Larval mortality rate | day<sup>-1</sup> |
| $\mu_P(T)$ | Pupal mortality rate | day<sup>-1</sup> |
| $\mu_A(T)$ | Active adult mortality rate | day<sup>-1</sup> |
| $\mu_r$ | Host-seeking mortality (adults) | day<sup>-1</sup> |
| $\mu_{A_d}$ | Diapausing adult mortality rate | day<sup>-1</sup> |
| $\sigma_L$ | Larval density-dependent mortality coefficient | day<sup>-1</sup> ind<sup>-1</sup> |
| $\delta(F)$ | Diapause entry rate | day<sup>-1</sup> |
| $\gamma(F)$ | Diapause termination rate | day<sup>-1</sup> |
| $T$ | Temperature | °C |
| $R$ | Rainfall | mm day<sup>-1</sup> |
| $F$ | Photoperiod | hours |

**Table 2.**
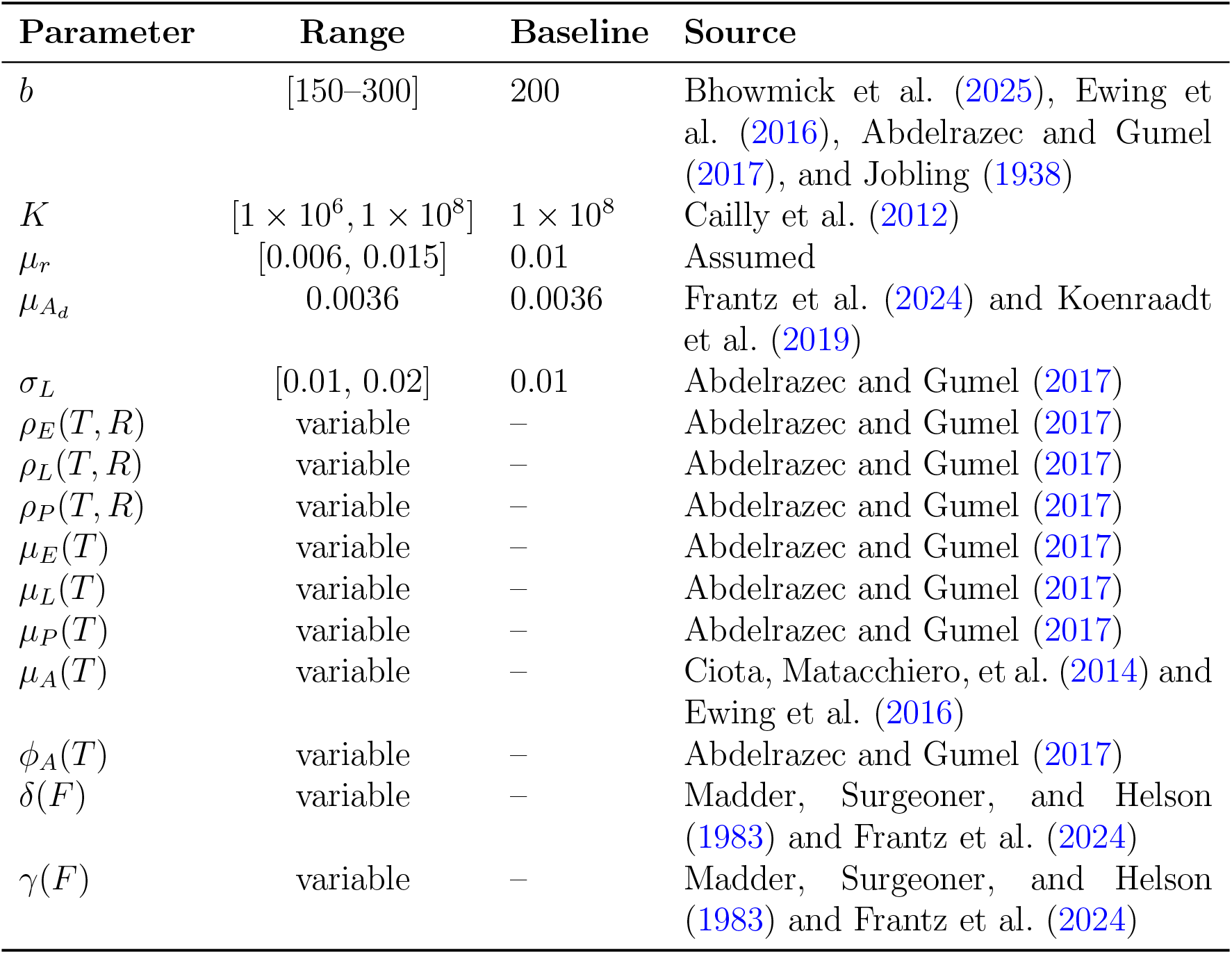
Model parameter values, units, and sources.

### 2.2 Data and Parameterization

#### 2.2.1 Environmental Drivers

The model incorporates three primary environmental drivers, temperature *T* (*t*), rainfall *R*(*t*), and photoperiod *F* (*t*), each influencing multiple stage-specific biological rates. Historical daily records for all three variables were obtained for the study location (Appendix A.3), averaged or sampled over multiple years to capture seasonal patterns. Where possible, smooth periodic functions were fitted to the data to provide continuous, bounded representations for use in simulations. Rainfall, due to its high day-to-day variability, was treated as a stochastic process. We describe the data sources, processing steps, and resulting functional forms for each driver.

Daily average temperature data from 2008–2023 (excluding 2012 due to missing values) were averaged across years to obtain a smooth seasonal profile. We then fit a periodic cosine function of the form

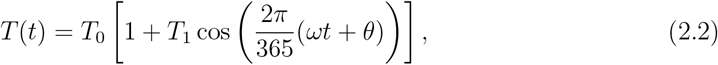

where *t* is the day of year, *T*_0_ is the mean temperature, *T*_1_ the relative amplitude, *ω* the frequency scaling, and *θ* the phase shift. This fitted seasonal curve (Figure 2a) captures the annual cycle and provides a continuous input for temperature-dependent development, reproduction, and mortality rates.

**Figure 2.**
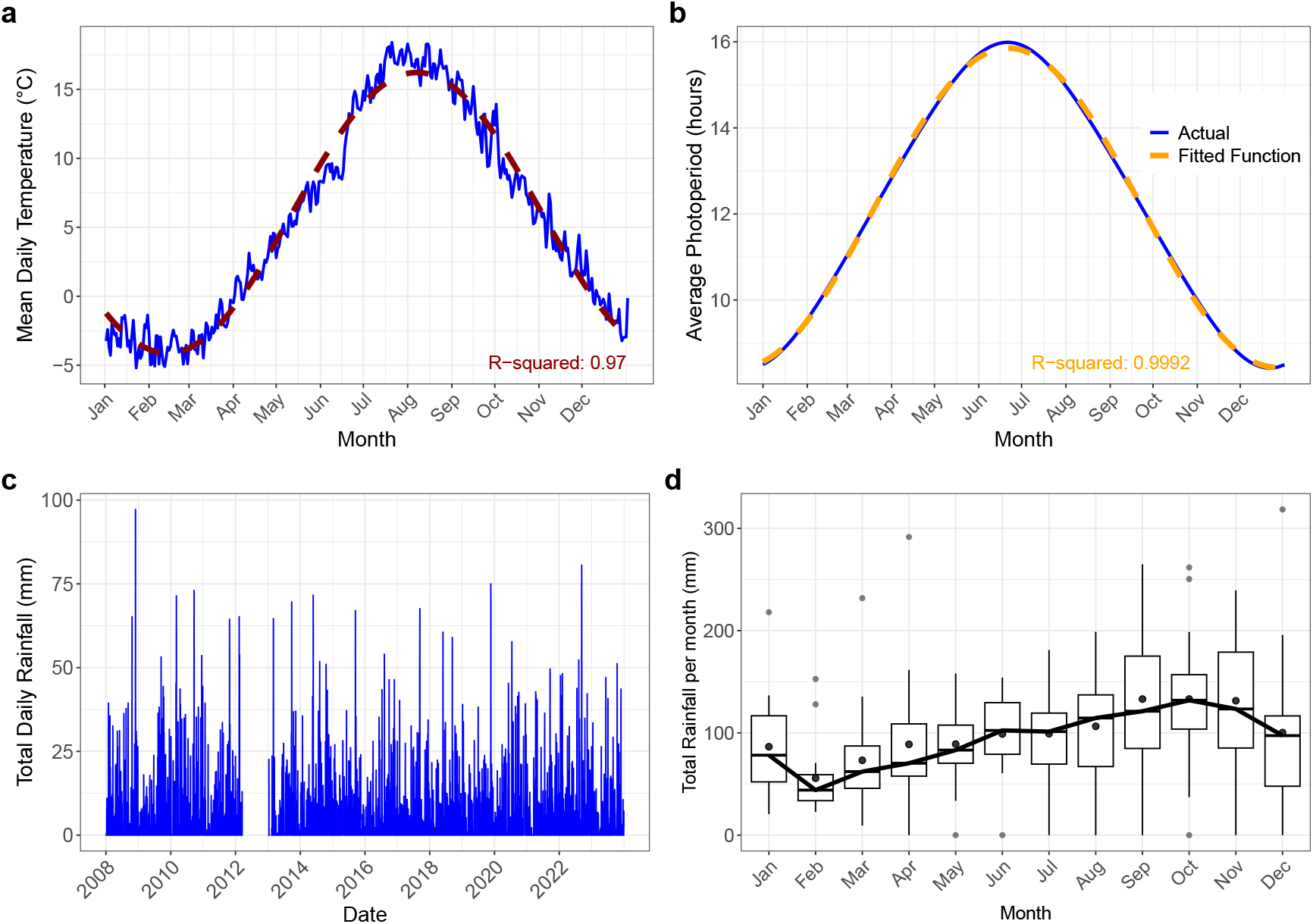
Environmental drivers used in the *Culex* population model. (a) Observed mean daily temperature (blue) and fitted periodic cosine function (red dashed line) for the study location, based on 2008–2023 records excluding 2012. (b) Observed mean daily photoperiod (blue) and fitted periodic cosine function (orange dashed line), based on 2008–2023 records. The fitted curve is used to drive diapause entry and termination. (c) Observed daily total rainfall across 2013–2023, illustrating strong day-to-day variability. (d) Monthly aggregated rainfall totals (boxplots across years) showing pronounced seasonality. Rainfall is treated as a stochastic driver by sampling daily values from the historical record within the same month, preserving seasonal distributions while introducing natural variability.

Daily total rainfall data exhibit high inter-day variability, making them unsuitable for fitting a smooth periodic function. Instead, rainfall is treated as a stochastic variable. To preserve observed seasonality while incorporating randomness, we applied a month-wise resampling procedure: for each simulated day, a rainfall value was randomly drawn (with replacement) from the same calendar month in the historical record (2013–2023). This approach maintains the empirical seasonal distribution of rainfall while reflecting the stochastic nature of precipitation events. The pronounced seasonality and short-term variability of rainfall are illustrated in Figure 2c&d, which shows both the daily fluctuations and monthly aggregated distribution.

Photoperiod, the number of daylight hours per day, regulates diapause induction and termination in adult mosquitoes. We computed the photoperiod *F* (*t*) programmatically in R using the `suncalc` package, which calculates sunrise and sunset times for a given latitude and longitude. The difference between sunrise and sunset times, computed via the `difftime` function, yields the daily photoperiod. Daily values were averaged over 2008–2023 to obtain a smooth seasonal profile, which was fit with a periodic cosine function:

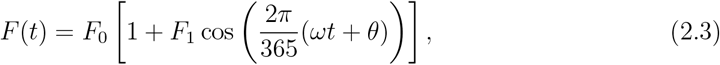

where parameters are defined analogously to the temperature function. This fitted curve (See Figure 2b) drives the diapause entry rate *δ*(*F*) and termination rate *γ*(*F*) in the model. Together, Figure 2 illustrate the seasonal patterns and variability of the environmental drivers used in the model, providing the empirical basis for the functional relationships and stochastic sampling schemes described in the following parameter function subsections.

### 2.3 Parameter Functions

The model incorporates temperature-, rainfall-, and photoperiod-dependent functions to represent key life-history processes of *Culex* mosquitoes. Parameterizations are either based on functional forms in previous studies or on empirical relationships from laboratory and field studies, with values drawn from literature where available. Where data were limited, parameters were estimated within biologically plausible ranges. The following subsections describe each functional form used in the model.

#### 2.3.1 Mortality rates

For juvenile mosquito stages (eggs *E*, larvae *L*, pupae *P*), temperature-dependent mortality rates follow the quadratic form used by Abdelrazec and Gumel (2017) for *Culex* in Ontario,

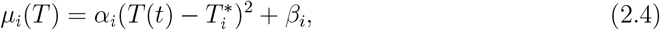

where *i* ∈ {*E, L, P*}, *T* (*t*) is temperature, 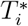 is the optimum temperature, and *α*_*i*_, *β*_*i*_ are fitted parameters.

Adult active-stage mortality is modeled using an exponential function fitted to laboratory data from Ciota, Matacchiero, et al. (2014), which was also used in Ewing et al. (2016),

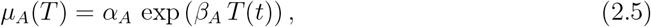

where *α*_*A*_ and *β*_*A*_ are fitted constants.

Estimating daily mortality during diapause, 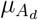 is challenging due to limited direct measurements. Studies suggest that 50–60% of diapausing adults survive winter (Koenraadt et al., 2019; Frantz et al., 2024). To approximate the duration of the winter period, we define winter in Newfoundland as November 15 to May 1 (Newfoundland and Labrado, 1998), lasting 167 days. Using the survival–mortality relationship from Frantz et al. (2024):

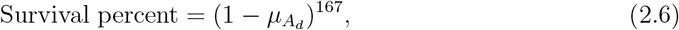

and assuming 55% survival, we computed 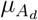 (see Table 2). Full calculation details are in the Appendix A.3.2.

#### 2.3.2 Oviposition rate

The adult oviposition rate *ϕ*_*A*_(*T*) was modeled as a temperature-dependent Gaussian function adapted from Abdelrazec and Gumel (2017),

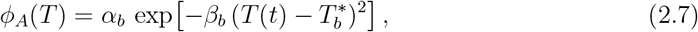

where *α*_*b*_ represents the peak oviposition rate, *β*_*b*_ controls the spread around the optimum, and 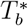 is the temperature at which oviposition is maximized. This functional form captures the observed nonlinear response of oviposition to temperature. In the cool-temperate Newfoundland context, breeding sites are rarely limiting due to permanent ponds and wetlands. Thus, oviposition is assumed to be driven primarily by temperature. This choice improves biological realism for the study region while avoiding unnecessary model complexity.

#### 2.3.3 Development rates

The maturation rate from stage *i* to the next stage depends on temperature and rainfall,

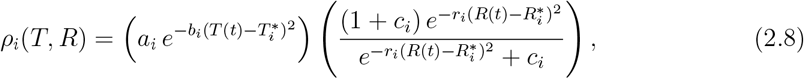

where *a*_*i*_ is the maximum rate, 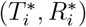 are the optimal temperature and rainfall, *b*_*i*_ and *r*_*i*_ control how sharply the rate declines away from these optima, and *c*_*i*_ controls rainfall saturation. This functional form is adapted from Abdelrazec and Gumel (2017), which was used to study *Culex* species in the Peel region of Ontario.

#### 2.3.4 Diapause Functions

Adult diapause dynamics are modeled using photoperiod-dependent functions fitted to experimental data from Madder, Surgeoner, and Helson (1983), based on field observations in summer 1979. For each observation date, corresponding daylight hours were calculated to match observed diapause rates with photoperiod.

Diapause induction is described by a logistic function (also used in Frantz et al. 2024):

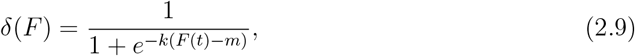

where *F* (*t*) is day length, *k* is the slope, and *m* is the midpoint threshold. Parameters were estimated in R using the nls() function, with fitted values reported in Table 4.

**Table 3.** Summary of parameter functions, biological processes, environmental drivers, and equation references.

| Function | Process | Driver(s) | Eq. |
| --- | --- | --- | --- |
| $\mu_i(T)$ | Mortality (juveniles $E, L, P$) | $T$ | 2.4 |
| $\mu_A(T)$ | Mortality (active adults) | $T$ | 2.5 |
| $\mu_{A_d}$ | Mortality (diapausing adults) | Winter duration | 2.6 |
| $\phi_A(T)$ | Oviposition rate | $T$ | 2.7 |
| $\rho_i(T, R)$ | Development ($E \rightarrow L \rightarrow P \rightarrow A$) | $T, R$ | 2.8 |
| $\delta(F)$ | Diapause induction | $F$ | 2.9 |
| $\gamma(F)$ | Diapause termination | $F$ | 2.10 |

**Table 4.**
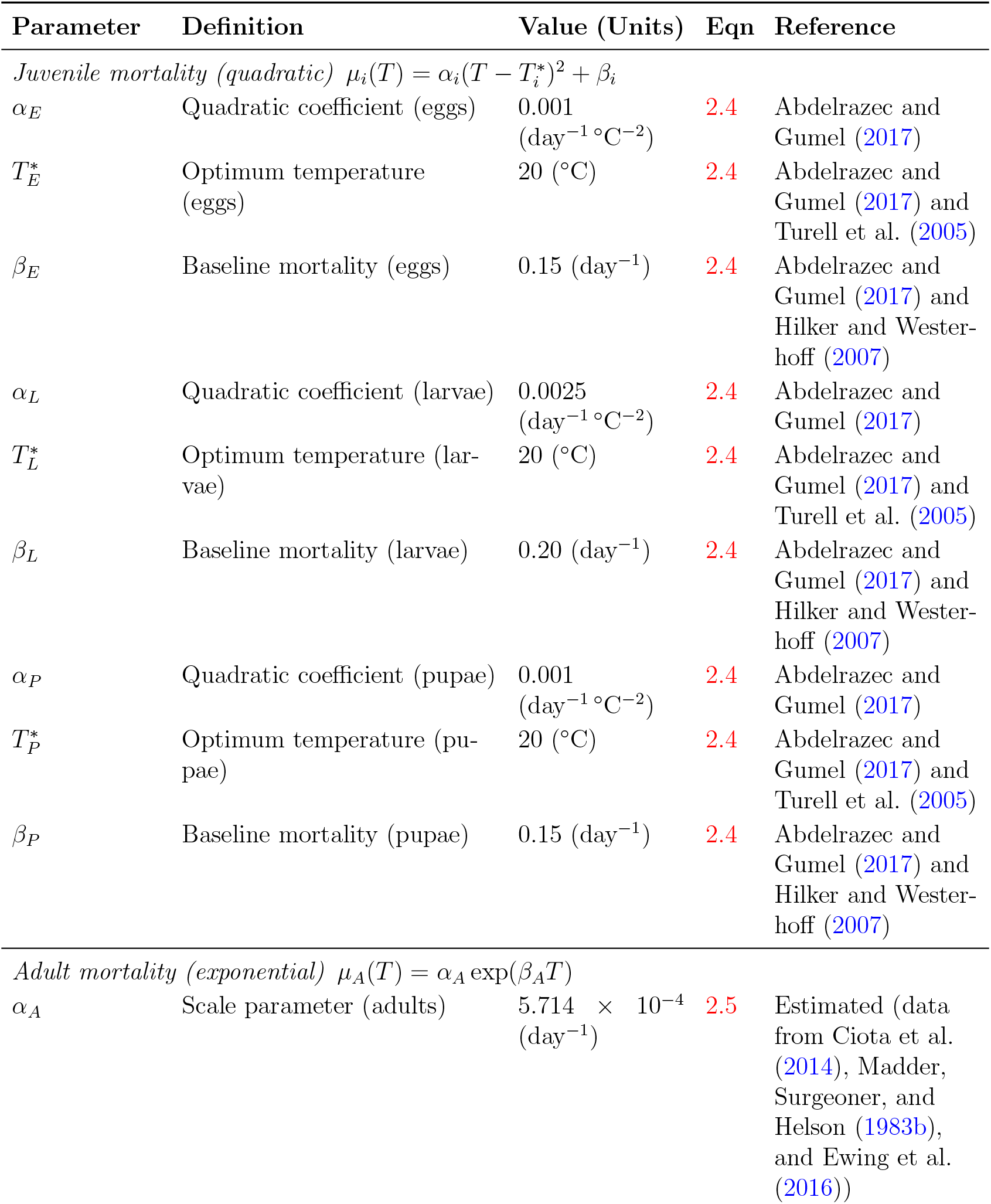
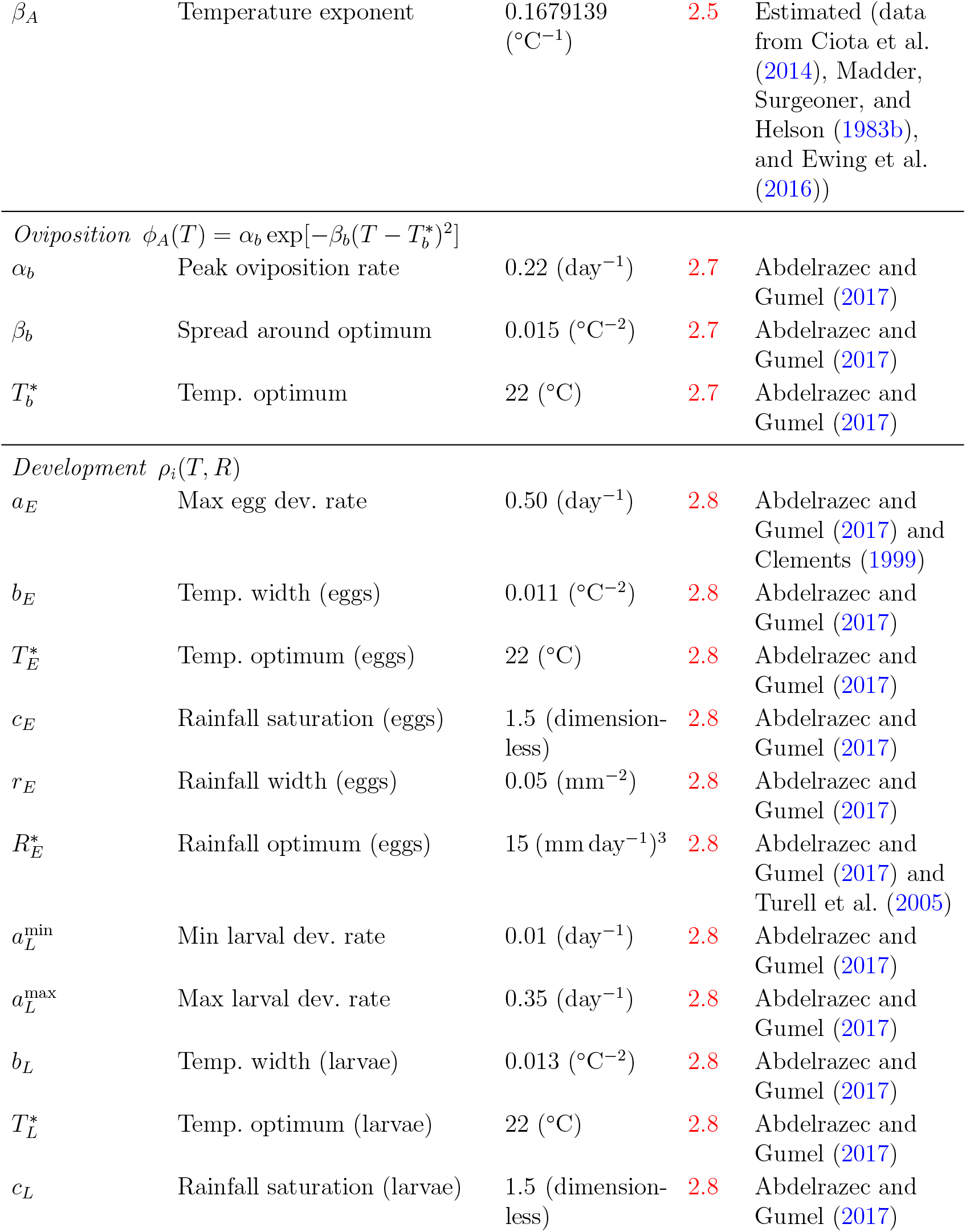
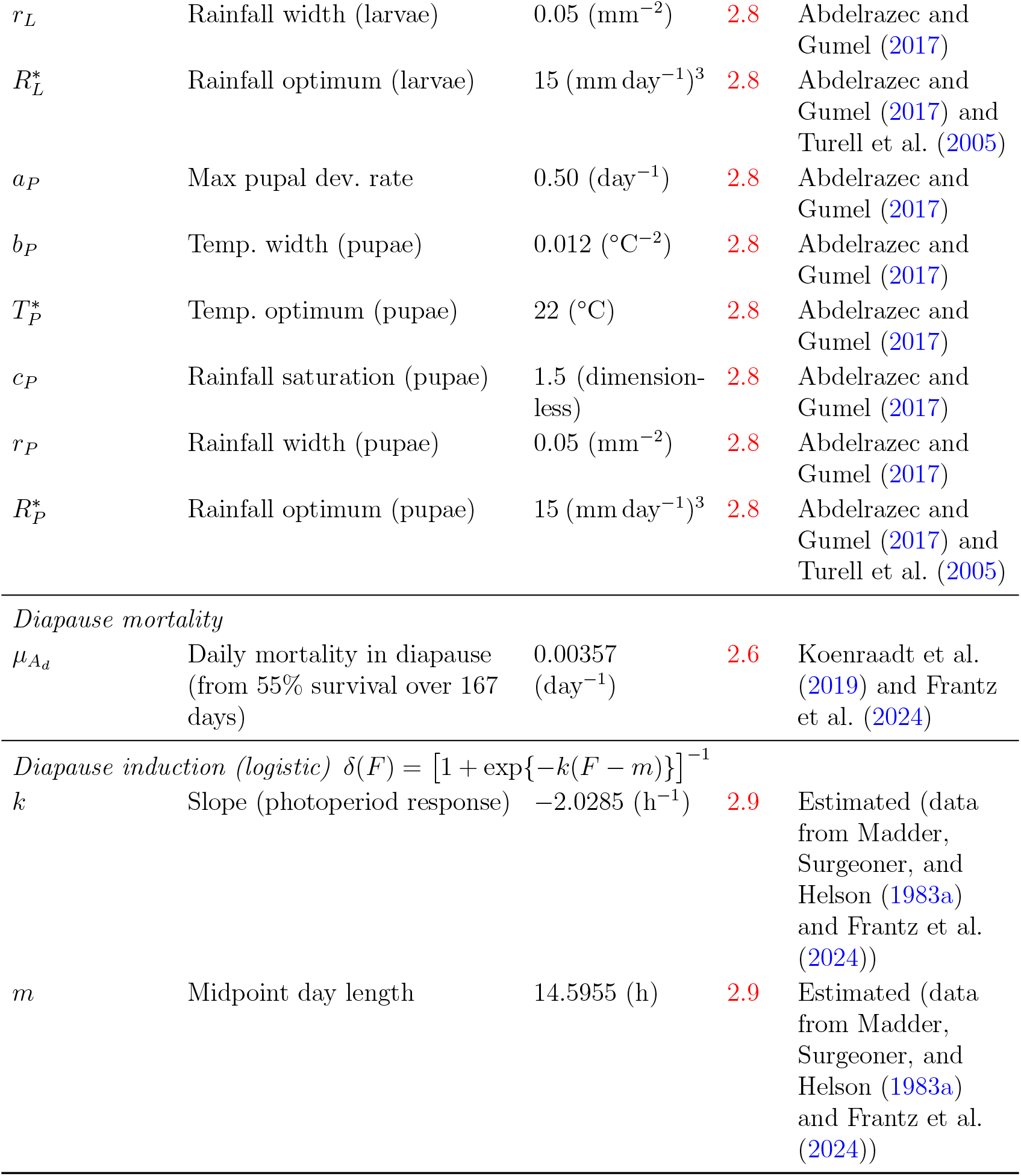
Fitted parameters used in the parameter functions.

Diapause termination is modeled as the complement of induction:

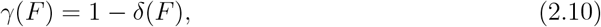

where *γ*(*F*) is the rate at which mosquitoes exit diapause. As day length increases, *γ*(*F*) rises, indicating a higher probability of mosquitoes becoming active; shorter day lengths reduce *γ*(*F*) as diapause activation dominates.

All parameter functions described above, along with the associated biological processes, environmental drivers, and corresponding equation numbers, are summarized in Table 3.

### 2.4 Numerical Implementation

The numerical solutions of the model system (A.1) were obtained using the `ode` function from the `deSolve` package in R (Soetaert, Petzoldt, and Setzer, 2010). The simulations were run over a five-year period with a daily time step (Δ*t* = 1 day) to match the temporal resolution of the input data. Parameterized functions, as described in Section 2.3, and constant parameters listed in Tables 2 and 4, were used for all analyses. Initial conditions for the simulations were assumed to be *E*(0) = 100, *L*(0) = 100, *P* (0) = 100, *A*(0) = 100, *A*_*d*_(0) = 100.

The numerical analysis aimed to examine how local environmental conditions, specifically temperature and rainfall, shape mosquito population dynamics and stage structure. This computational framework provided the basis for the scenario experiments.

### 2.5 Simulation Scenarios

To evaluate the model under varying climatic conditions, simulations were conducted for a set of predefined scenarios representing baseline and perturbed environments. The baseline scenario used observed seasonal cycles of temperature and photoperiod, together with stochastic rainfall sampling that preserved the historical seasonal structure. Perturbation scenarios were designed to test the effects of altered temperature and rainfall regimes, both individually and in combination.

Temperature perturbations were implemented as uniform shifts of the baseline seasonal cycle, ranging from +1°C to +5°C. Rainfall perturbations were introduced by modifying the sampled daily rainfall time series to represent ‘dry,’ ‘wet,’ and ‘erratic’ conditions, whereby daily totals were reduced by a fixed percentage, increased by the same magnitude, or adjusted using randomized multiplicative factors to simulate greater variability. In both cases, photoperiod and the non-perturbed climate variable were retained from the observed historical records to isolate the effect of the perturbation under consideration. To examine potential interactions, combined scenarios were simulated in which each temperature shift was paired with each rainfall regime, enabling assessment of additive and nonlinear effects arising from concurrent changes in temperature and rainfall.

Each scenario was simulated for 100 independent replicates to capture stochastic variability arising from the rainfall sampling procedure. Replicate outputs were stored in structured data frames for subsequent statistical analysis, including computation of mean trajectories and variability measures. Further details on the computation of summary statistics, prediction intervals, and visualization of results are provided in Appendix A.2.

We summarized model dynamics using biologically relevant metrics, stage-specific dynamics, mean adult abundance, peak abundance, and duration of the active season (defined as the period when temperature exceeded 10°C leading to active adult population surpassing the diapausing population). These measures provide ecologically meaningful insights into seasonal mosquito activity and potential impacts on disease transmission risk.

## 3 Results

Model simulations revealed clear seasonal patterns in *Culex* population dynamics, shaped by interactions between temperature, rainfall, and photoperiod. We first describe stage-structure dynamics under observed climatic conditions, followed by comparisons across temperature and rainfall perturbation scenarios. We then present the outcomes of the combined effect of temperature shifts and rainfall scenarios on the adult abundance and peak timing. Together, these results provide an integrated view of how environmental variability, climate warming and life-history processes shape mosquito population dynamics and seasonal activity.

### 3.1 Seasonal Dynamics and Variability Across Life Stages

Figure 3 shows simulated trajectories of *Culex* populations across all life stages during years 3–5 under 100 stochastic rainfall realizations. Eggs (E), larvae (L), and pupae (P) exhibited sharp summer peaks followed by steep declines in fall. These early stages displayed the greatest variability, with eggs reaching the highest abundances and widest uncertainty ranges, highlighting strong environmental sensitivity. Adults (A) also peaked annually but with narrower prediction intervals, indicating more consistent dynamics across simulations. This stability likely reflects their longer lifespan and the smoothing of variability carried over from earlier stages. Diapausing adults (*A*_*d*_) persisted year-round, accumulating in late summer and fall before stabilizing through winter. Although their overall abundance varied less, greater uncertainty occurred in the timing of transitions into and out of diapause, consistent with photoperiod and temperature cues rather than rainfall as primary drivers.

**Figure 3.**
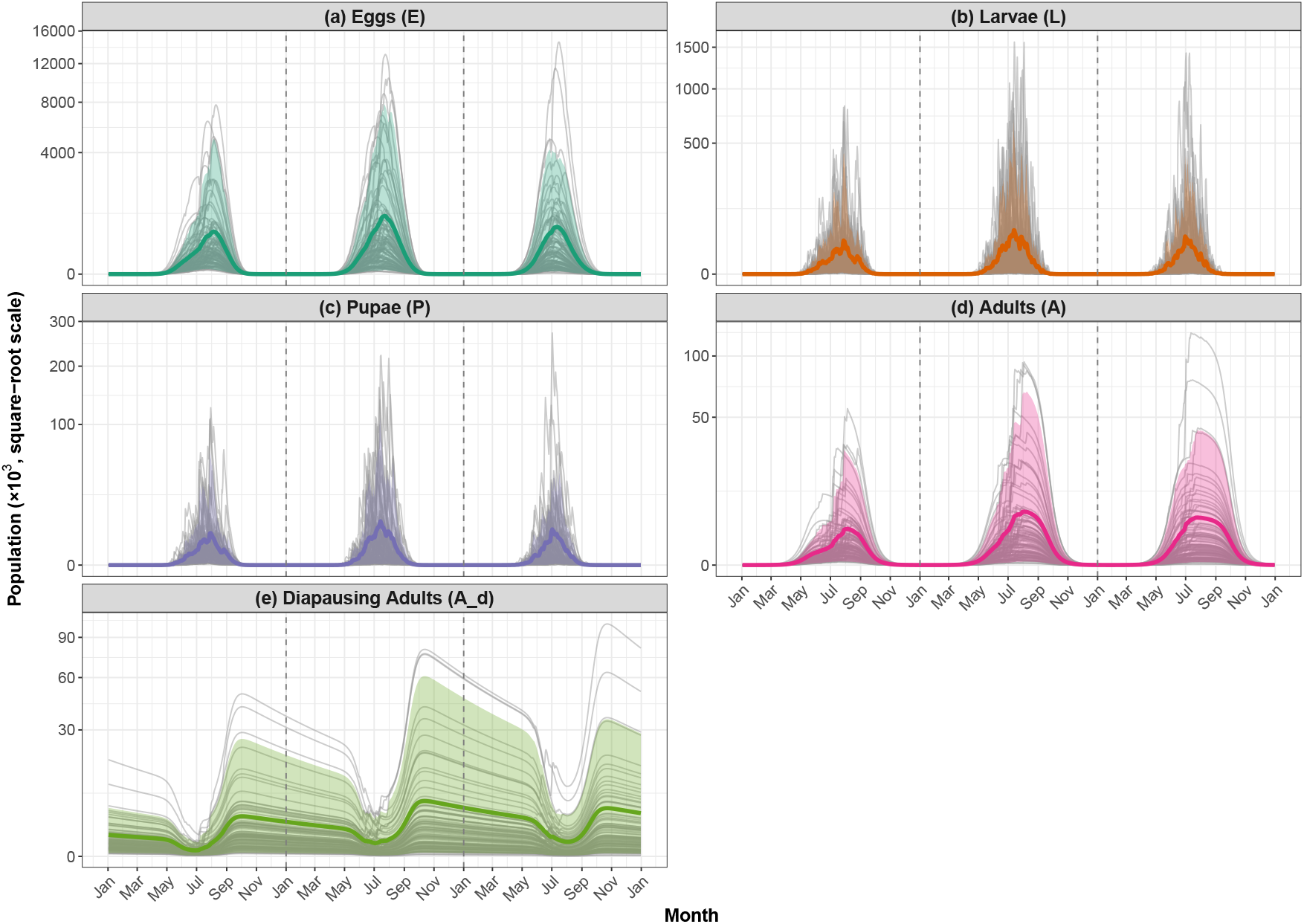
Stage-specific population trajectories of *Culex* mosquitoes over simulation years 3 to 5 under 100 stochastic rainfall realizations. Each panel corresponds to a distinct life stage, eggs (E), (b) larvae (L), (c) pupae (P), (d) active adults (A), and (e) diapausing adults (*A*_*d*_). Gray lines represent individual simulation runs, colored bold lines indicate the mean population trajectory, shaded bands show 95% prediction intervals. The mean and prediction intervals highlight the typical seasonal dynamics and quantify the uncertainty arising from stochastic rainfall, offering insight into the variability of the abundance of each stage. Only simulation years 3 to 5 are shown to minimize the influence of assumed initial state values and capture long-term dynamics. Rainfall was sampled with replacement by calendar month to preserve seasonal structure, while temperature and photoperiod were held deterministic across runs. Vertical dashed lines mark the boundaries between years 3, 4, and 5.

### 3.2 Temperature-driven shifts in adult abundance

Simulations revealed strong temperature effects on mosquito dynamics. At baseline (+0°C), adult populations remained low and largely restricted to summer months. Warming amplified both the magnitude and duration of seasonal peaks, with abundance increasing by several orders of magnitude at +2°C and +3°C (Figure 4). At higher warming (+4°C and +5°C), extremely dense summer populations persisted, although populations still collapsed during winter. These results indicate that warming both elevates abundance and extends the seasonal window of mosquito activity.

**Figure 4.**
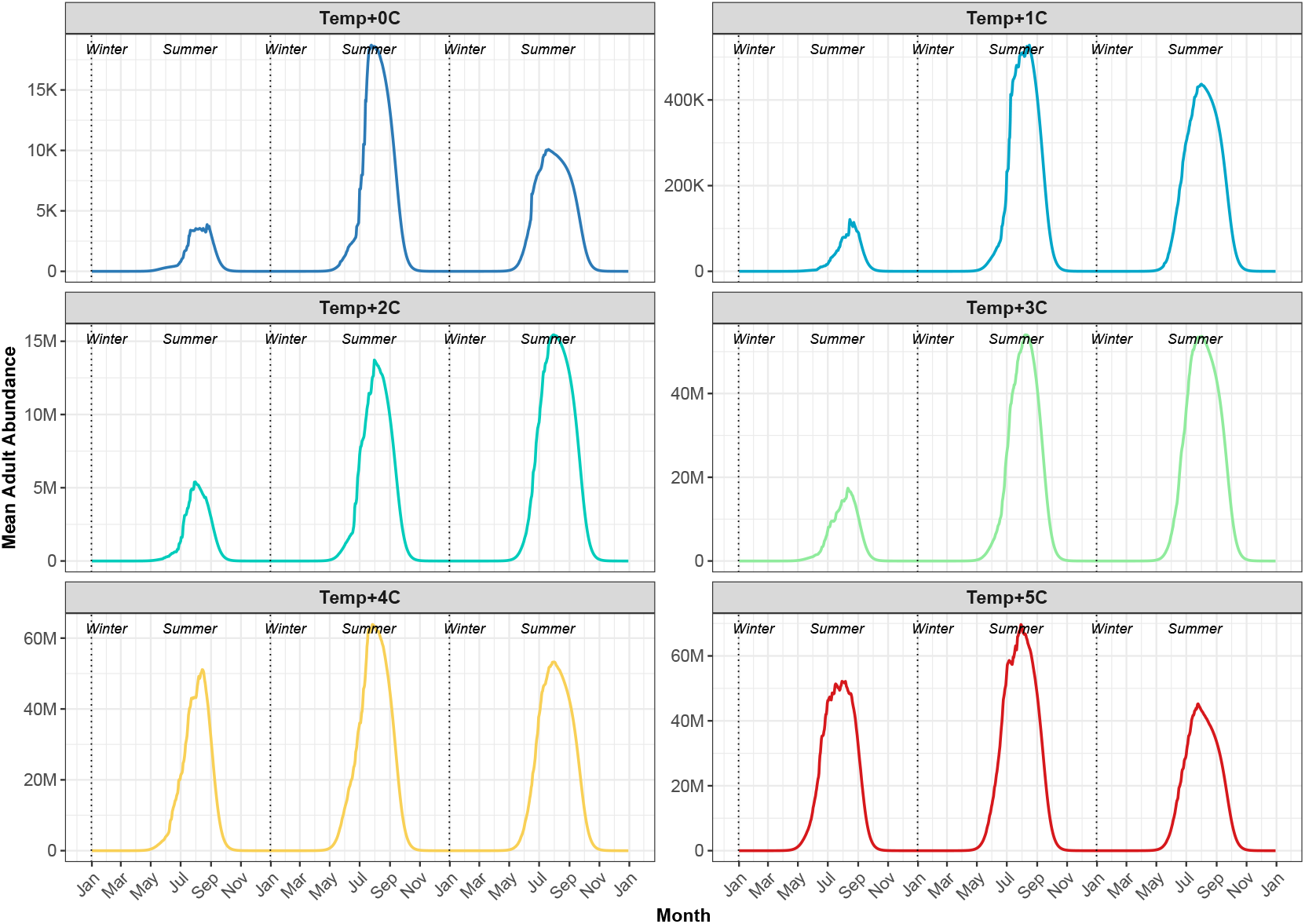
Mean adult mosquito abundance trajectories across 100 replicate simulations for each temperature shift. Each replicate used a different stochastic rainfall realization. Temperature shifts range from baseline (*Temp* + 0^*°*^)C to +5° C. Each line represents the average daily adult abundance across replicates. Vertical dashed lines indicate year boundaries over the five-year simulation period. Seasonal population dynamics emerge and become increasingly pronounced and sustained with warming.

Total adult abundance over five years highlighted the nonlinear response to temperature (Figure 5). Abundance increased nearly four orders of magnitude between baseline and +2°C, but plateaued under stronger warming. Stochastic variability across rainfall replicates was greatest at intermediate warming (+1°C to +2°C), reflecting strong interactions between rainfall and temperature, whereas variability diminished at higher warming, where consistently high densities were maintained regardless of rainfall.

**Figure 5.**
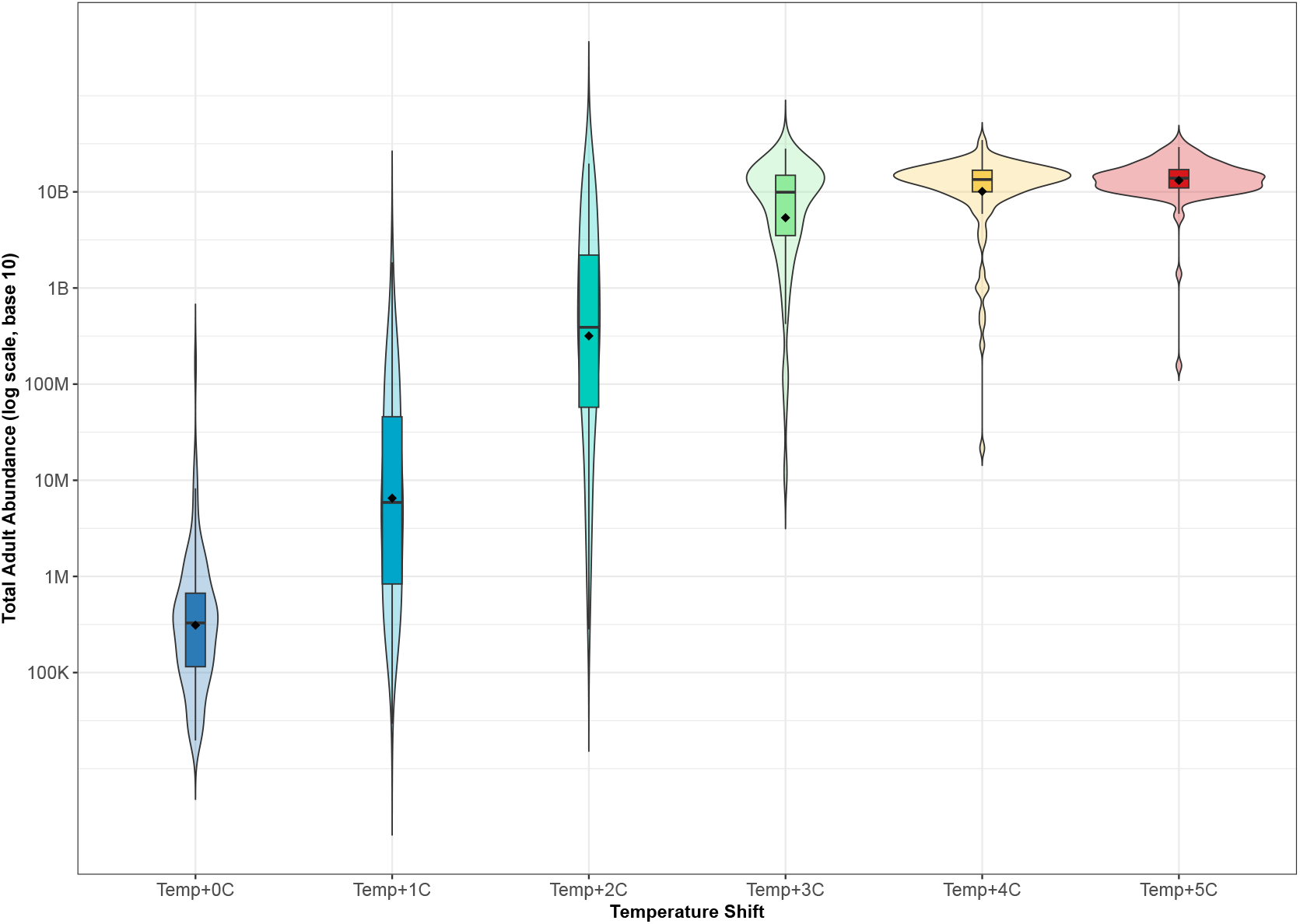
Distribution of total adult mosquito abundance across five years under temperature shifts. Black dots indicate the mean abundance across rainfall realizations. Total abundance rose sharply from baseline to +2°C, then leveled off at consistently high values under +3°C to +5°C warming.

### 3.3 Active Days Across Temperature Shifts

The analysis revealed a pronounced effect of warming on the seasonal activity of *Culex* mosquitoes. Figure 6a shows that higher temperature shift advanced the onset of activity into early spring (March–April) and extended it later into the fall (October-November), thereby increasing the duration of the active season. This seasonal expansion was most pronounced under the +5°C shift, which maintained near-maximal activity from May through September. Figure 6b summarizes the cumulative effect of warming on mosquito activity, showing that average active days increased progressively with temperature shifts, ranging from a 9.1% increase under +1°C to a 42.9% increase under +5°C. Together, these results highlight both the shift in seasonal timing and the overall intensification of mosquito activity expected under future warming scenarios.

**Figure 6.**
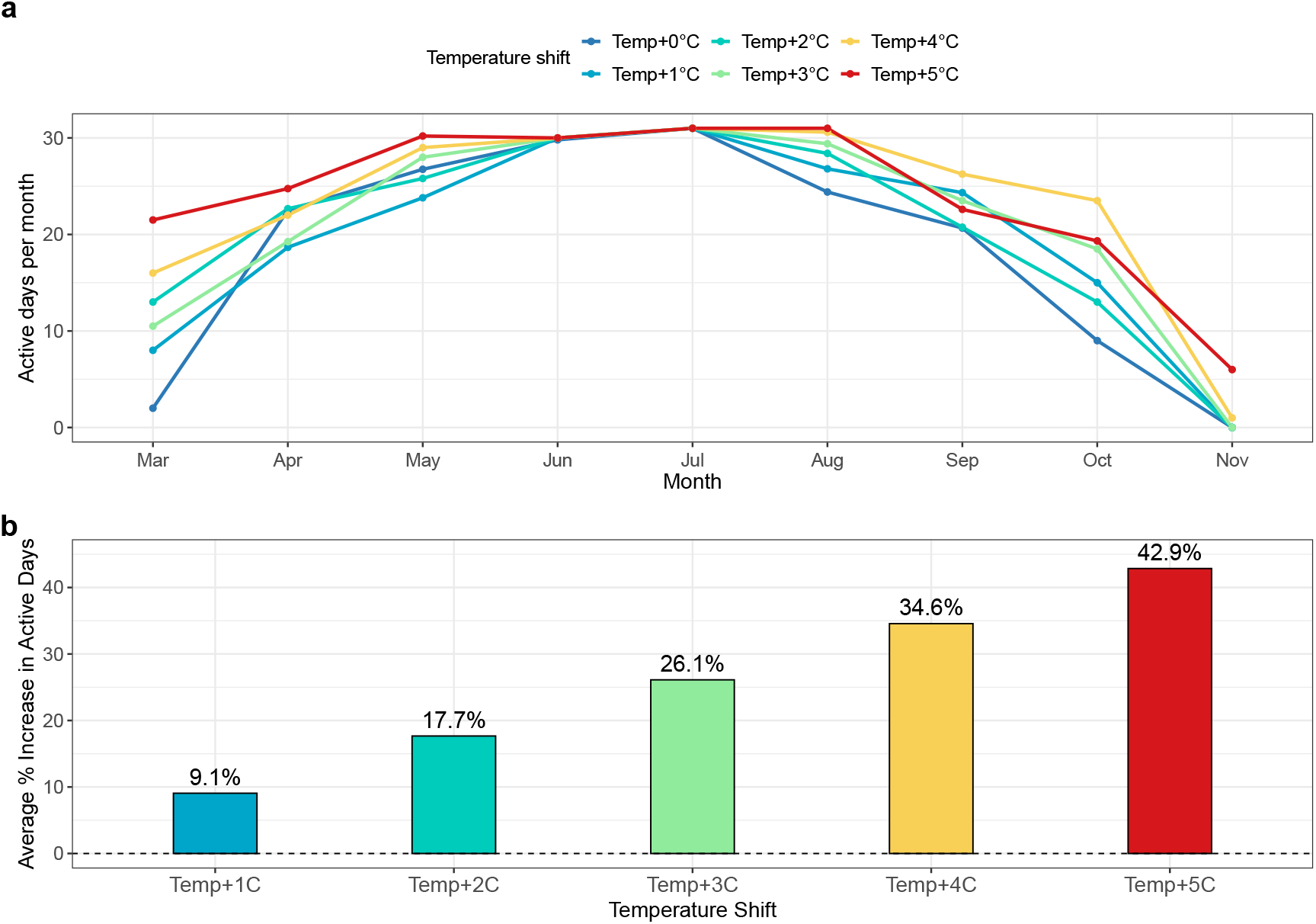
Effect of temperature shifts on mosquito activity. (a) Monthly active days under baseline and warming scenarios, showing that warming extends mosquito activity into early spring and late fall. (b) Average percentage increase in active days relative to the baseline, summarizing the cumulative effect of warming across temperature shifts. Together, these panels highlight both seasonal shifts and the overall amplification of mosquito activity under climate warming.

### 3.4 Rainfall-driven variation in mosquito abundance

Rainfall strongly influenced mosquito dynamics, with clear contrasts across scenarios (Figure 7). Daily rainfall patterns showed persistently low inputs in the dry scenario, consistently elevated rainfall in the wet scenario, and the erratic scenario yielded intermediate outcomes, with variability comparable to or slightly higher than baseline, but without the extreme amplification observed under wet conditions (Figure 7b). These differences translated into seasonal rainfall totals, with the wet scenario yielding the greatest accumulations, the dry scenario the least, and the baseline and erratic cases falling in between (Figure 7c).

**Figure 7.**
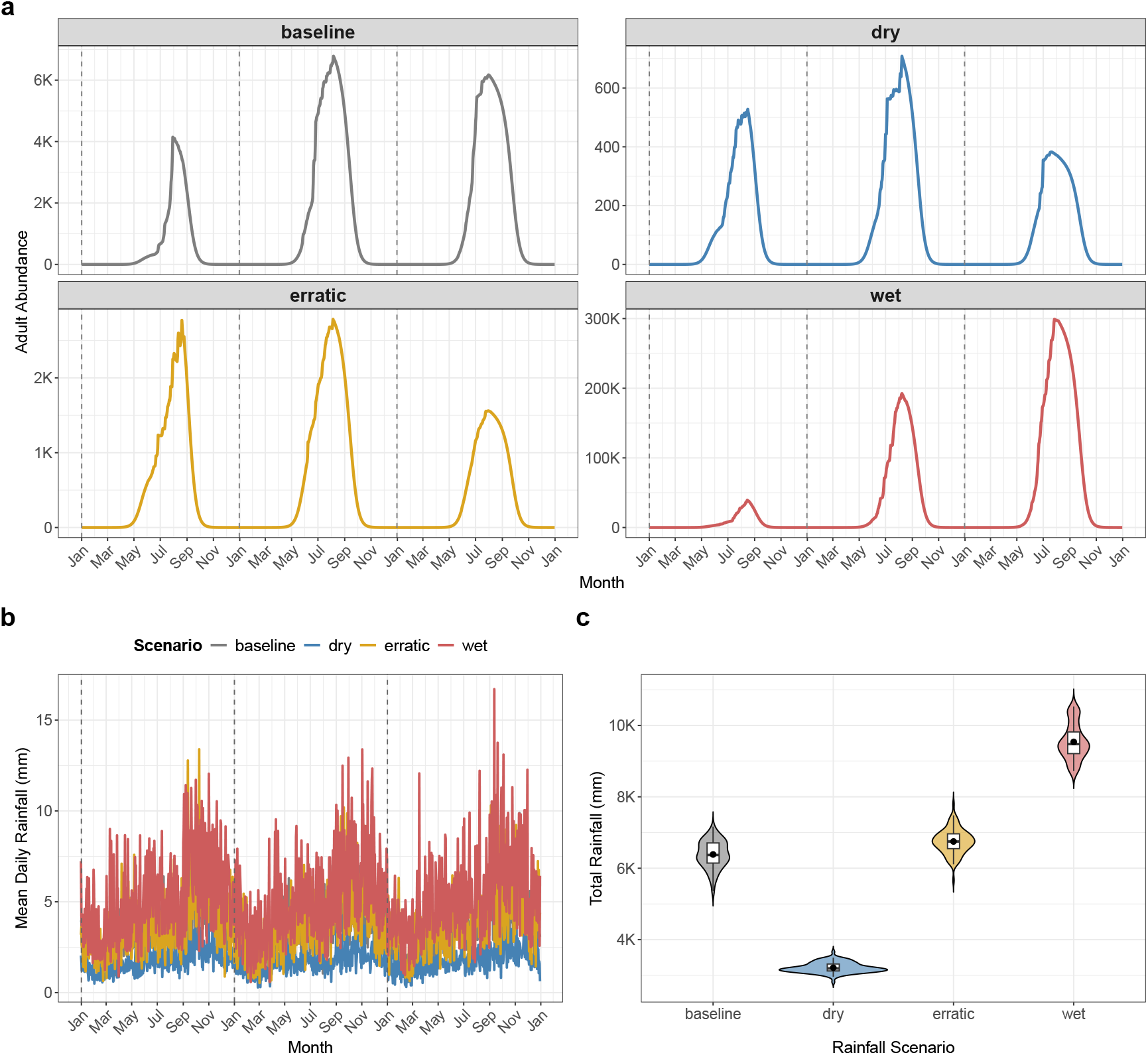
Rainfall scenarios and their effects on mosquito abundance. (a) Mean adult mosquito abundance under baseline, dry, erratic, and wet rainfall scenarios, showing suppressed populations under dry conditions and extreme amplification under wet conditions. (b) Mean daily rainfall across scenarios, highlighting the highest variability and magnitude under wet conditions and persistently low inputs under dry conditions. (c) Total seasonal rainfall distributions, illustrating clear contrasts across scenarios, with wet conditions yielding the highest accumulation and dry conditions the lowest. Grey dashed vertical lines indicate the start of each simulation year. Together, these results demonstrate that rainfall strongly influences both mosquito abundance and their temporal stability, amplifying abundance under wet conditions while constraining it under drought.

Mosquito abundance was strongly linked to rainfall availability (Figure 7a). Under baseline conditions, populations exhibited pronounced seasonal peaks but remained moderate overall. The dry scenario strongly suppressed abundance, producing consistently low peaks, whereas the wet scenario drove explosive growth, with summer populations far exceeding baseline levels. The erratic scenario generated intermediate outcomes, with peak sizes varying substantially across seasons. Together, these results show that rainfall regulates both the magnitude and variability of mosquito populations, amplifying abundance under wet conditions while sharply limiting it under drought.

To further quantify these differences, we compared distributions of total and peak adult abundance across scenarios (Figure 8). The patterns were consistent: the wet scenario produced the largest populations, the dry scenario sharply constrained them, and the erratic scenario yielded intermediate values. These comparisons highlight that rainfall not only shapes seasonal dynamics but also sets the overall scale of mosquito population size.

**Figure 8.**
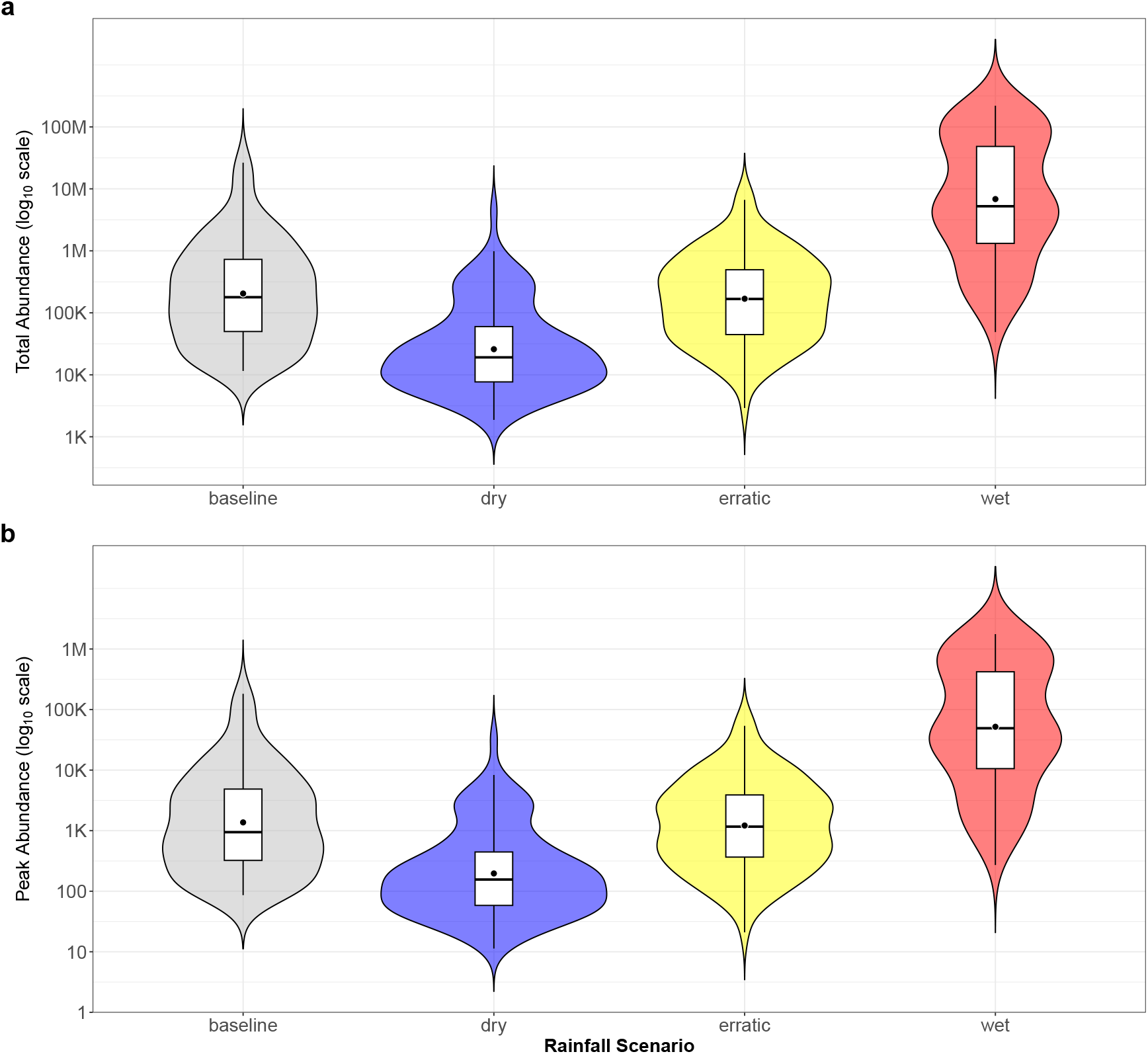
Distributions of (a) total adult mosquito abundance and (b) peak adult mosquito abundance across rainfall scenarios. Violin plots show the full distribution, with boxplots indicating the interquartile range and median, and dots marking the mean. The wet scenario produced the largest populations, the dry scenario the lowest, and the erratic scenario yielded intermediate outcomes close to the baseline.

### 3.5 Combined effects of temperature and rainfall on mosquito abundance

Temperature and rainfall interacted strongly to shape mosquito dynamics (Figure 9). Seasonal trajectories (Figure 9a) showed that under dry conditions, mosquito populations were restricted to low summer peaks. Warming amplified the magnitude of these peaks, with +2°C and +4°C shifts producing progressively larger summer populations across all rainfall scenarios. Rainfall modulated this response, dry conditions strongly suppressed abundance regardless of warming, while wet conditions drove explosive growth, with summer populations exceeding those observed under baseline rainfall. Erratic rainfall produced intermediate outcomes.

**Figure 9.**
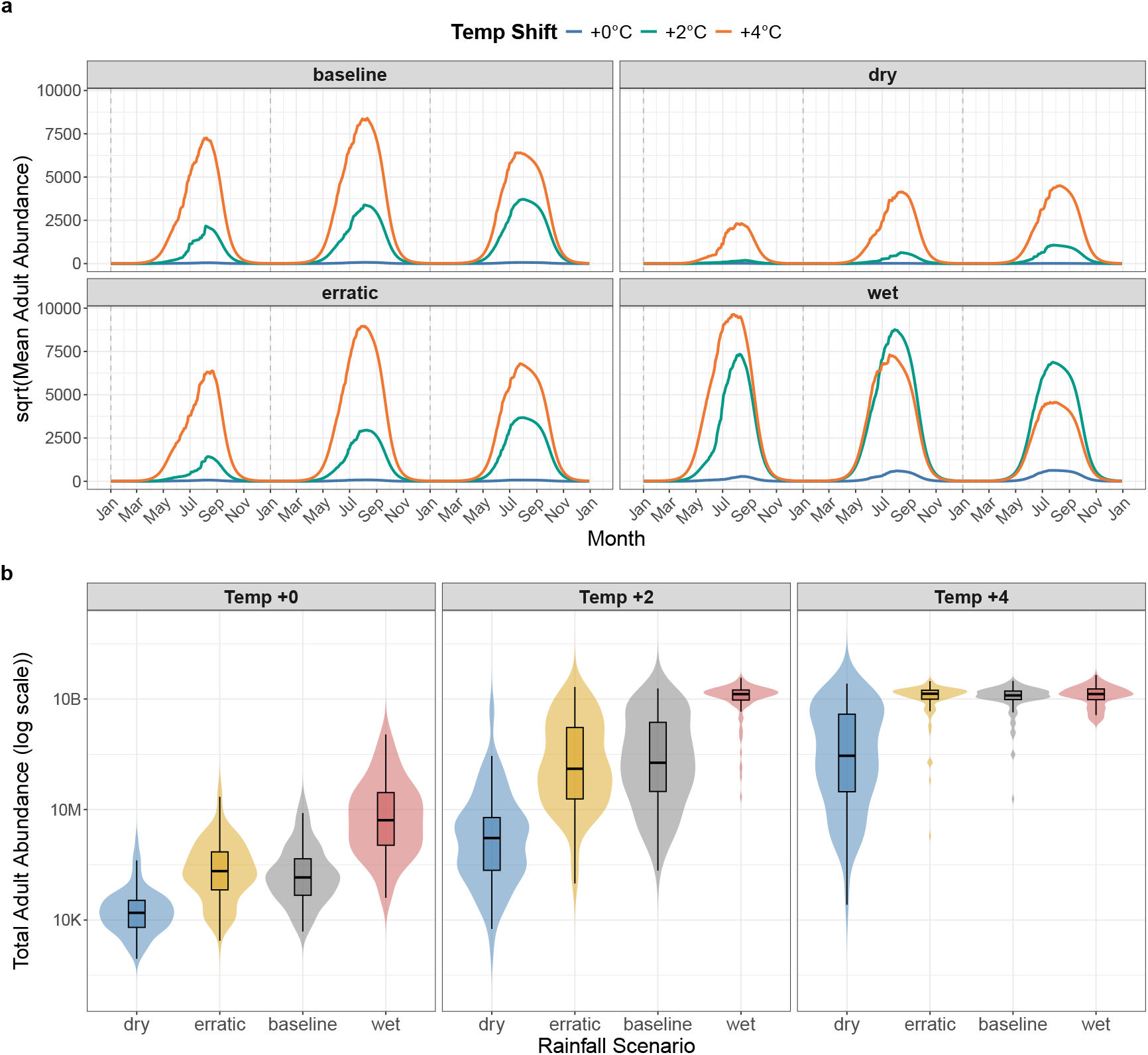
Combined effects of temperature and rainfall scenarios on mosquito abundance. (**a**) Seasonal trajectories of mean adult abundance under baseline, dry, erratic, and wet rainfall scenarios for three temperature shifts (+0°C, +2°C, +4°C). (**b**) Distributions of total adult abundance (log scale) across rainfall scenarios at each temperature shift.

Aggregated outcomes (Figure 9b) reinforced these patterns. Total adult abundance increased nonlinearly with warming, rising steeply from baseline to +2°C but plateauing at higher warming levels. Differences among rainfall scenarios were most pronounced under low and intermediate warming, dry conditions consistently yielded the lowest totals, wet conditions the highest, and erratic and baseline scenarios fell in between. At +4°C, abundances converged across scenarios, indicating that high temperatures can overwhelm the moderating influence of rainfall variability. This pattern suggests that at lower warming levels, rainfall differences play a stronger role in shaping mosquito dynamics, but under more extreme warming, populations converge toward high densities.

## 4 Discussion

### 4.1 Seasonal Trends and Uncertainty

Our results reveal strong stage-specific responses of *Culex* populations to environmental variability (as seen in Figure 3). Immature stages (eggs, larvae, and pupae) peaked sharply in summer with high variability across simulations, underscoring their strong rainfall sensitivity and the potential for moderate rainfall shifts to amplify seasonal population growth. In contrast, adults exhibited more stable dynamics, likely buffered by longer lifespans and demographic averaging, making them a more predictable and effective target for control. Diapausing adults followed a distinct seasonal rhythm, accumulating in late summer and persisting through winter, with transitions driven largely by photoperiod and temperature rather than rainfall. This stage is critical for overwintering and long-term persistence, though uncertainty in timing highlights variability in environmental cueing. Overall, the findings show how rainfall amplifies early-stage variability while adults remain comparatively stable, and how diapause dynamics underscore the importance of seasonal drivers.

### 4.2 Implications of temperature-driven shifts in adult abundance and active season

Our simulations demonstrate that mosquito populations are highly sensitive to warming, with strong nonlinearities in both abundance and seasonal persistence. At baseline conditions (+0°C), adult populations were restricted to short-lived summer peaks (see Figure 4), consistent with observations in temperate regions where overwintering constrains vector activity. In contrast, even modest warming (+2°C to +3°C) produced dramatic increases in population size and extended the active season (see Figures 5 and 6). With modest warming expected over the coming decades (+2°C to +3°C) (IPCC, 2021), regions currently characterized by short mosquito activity may experience markedly longer periods of mosquito activity, amplifying the potential for disease spread. The nonlinear responses we observed align with previous studies showing threshold effects of temperature on mosquito development and survival (Mordecai et al., 2019; Reinhold, Lazzari, and Lahondère, 2018). Notably, stochastic rainfall introduced the greatest variability at intermediate warming levels, indicating that climate interactions may generate unpredictable dynamics near ecological thresholds. By contrast, at higher warming (+4°C to +5°C), populations stabilized at consistently high abundance (see Figure 5). This plateauing reflects the influence of the fixed environmental carrying capacity (*K* = 10^8^) used in the model, which was chosen to ensure numerical stability and comparability across scenarios. In extreme cases (e.g., +5°C under wet conditions), mosquito abundance approached or reached *K*, underscoring the strong amplifying effects of warming and rainfall. These outputs should therefore be interpreted as the model saturating at its technical ceiling, not as evidence of a true ecological bound. Future work could incorporate climate- or resource-dependent carrying capacities to capture more realistic upper limits on mosquito abundance. Such patterns nonetheless highlight the potential for both sharp, nonlinear increases in vector abundance and reduced interannual variability under continued warming.

From a public health perspective, these findings underscore the dual risks posed by climate warming: elevated mosquito abundance and a widening seasonal window of activity. Extended activity into early spring and late fall increases overlap with human exposure periods and may facilitate pathogen establishment in areas where transmission is currently constrained. Our results therefore support proactive intervention strategies, including enhanced surveillance, targeted vector control, and early-warning systems in regions projected to warm by 2–3°C.

### 4.3 Implications of rainfall-driven variation in mosquito abundance

Our results show that rainfall strongly shapes mosquito population dynamics, influencing both abundance and variability. Abundance was strongly linked to rainfall availability: dry conditions suppressed populations, wet conditions produced explosive growth, and the erratic scenario generated intermediate outcomes. These patterns are consistent with field evidence that rainfall governs oviposition success, larval habitat availability, and juvenile survival (Parham and Michael, 2009; Yang et al., 2009), emphasizing the potential for rainfall anomalies to alter both the magnitude and predictability of outbreaks.

The amplification of abundance under wet conditions highlights the risk that future increases in extreme rainfall events could intensify mosquito proliferation, while drought may constrain populations but still sustain breeding in residual habitats. Variability under erratic conditions underscores the challenge of forecasting mosquito risk in climates where rainfall is unpredictable, consistent with observations in semi-arid and monsoon regions (Githeko et al., 2000; Abbasi, 2025; Ryan et al., 2019; Morin, Comrie, and Ernst, 2013). These findings suggest that rainfall variability is important in determining mosquito risk. Incorporating variability of rainfall into predictive models and surveillance systems will be essential under climate change scenarios that project intensification of wet and dry extremes.

### 4.4 Implications of temperature–rainfall interactions for mosquito dynamics

Our results demonstrate that temperature and rainfall interact nonlinearly to shape mosquito populations. Moderate warming (+2°C) magnified the influence of rainfall, widening the differences in mosquito abundance among scenarios. Dry conditions suppressed populations, wet conditions produced large increases in abundance, and erratic rainfall yielded intermediate but variable outcomes. At higher warming level (+4°C), abundance converged toward consistently high densities, suggesting that extreme temperatures can diminish the moderating influence of rainfall variability. These findings highlight that climate variability are critical for predicting mosquito responses under climate change.

Ecologically, these patterns emphasize the dual role of temperature and rainfall in regulating mosquito populations. While warming extends the active season and raises baseline abundance, rainfall determines whether populations remain suppressed, fluctuate unpredictably, or become explosively amplified. This interaction suggest that climate variability must be incorporated into risk assessment. The convergence of abundances under higher warming underscores the risk of consistently elevated mosquito densities, suggesting that even regions with historically unfavourable rainfall may face heightened vector-borne disease risk under future climate change.

A global sensitivity analysis of constant parameters (See Appendix A.4, Figure 10) shows that the number of eggs laid per oviposition (*b*) and adult diapause mortality (*µ*_*Ad*_) had the strongest influence on abundance, while other parameters had weaker effects. This shows that climate drivers remain the dominant sources of variability, underscoring the importance of explicitly incorporating temperature and rainfall dynamics in models of mosquito populations.

**Figure 10.**
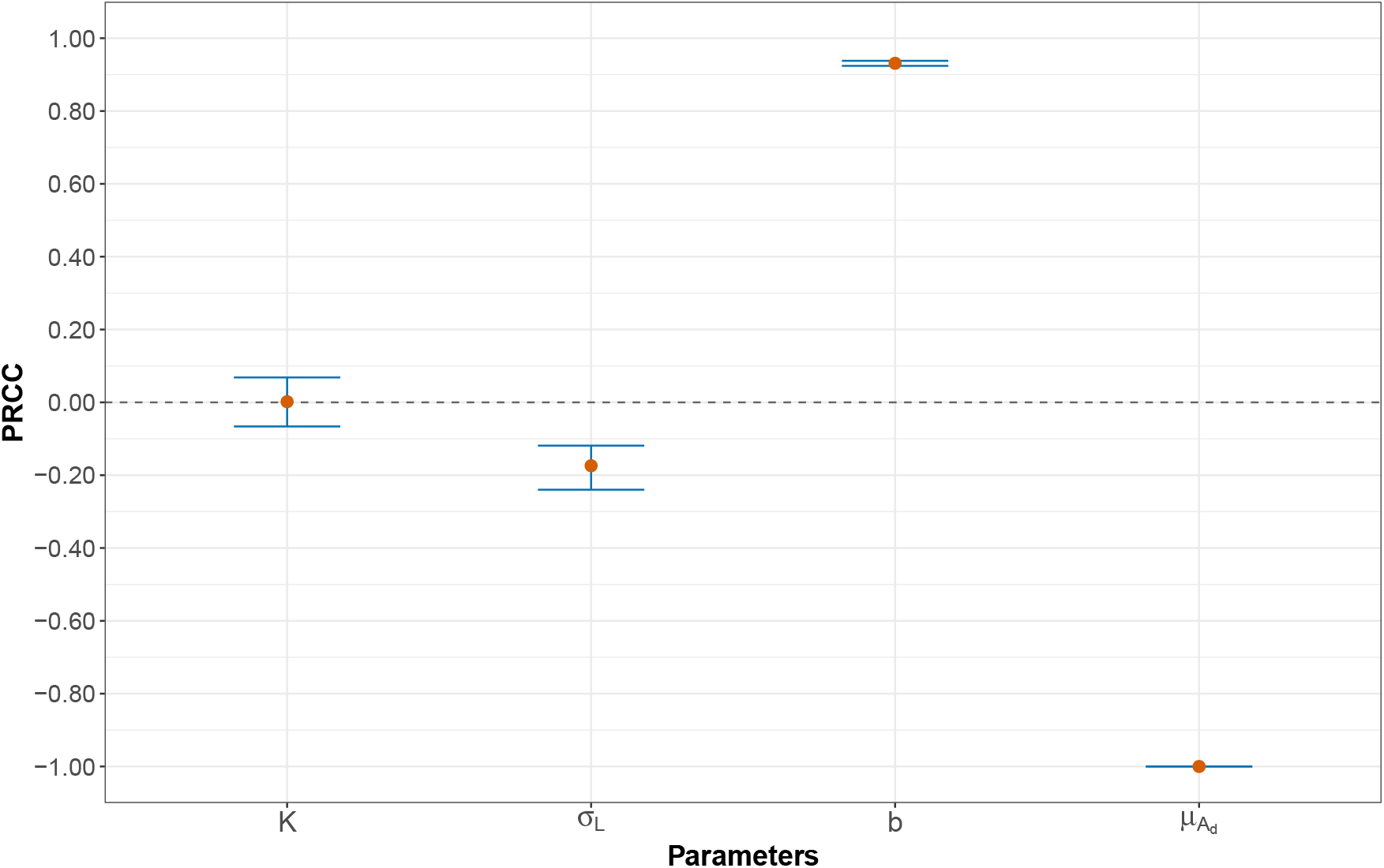
Global sensitivity analysis of constant parameters using Latin hypercube sampling (LHS) and partial rank correlation coefficients (PRCCs). Parameters are ordered along the x-axis by their absolute PRCC values. Points indicate mean PRCC values, with error bars showing 95% confidence intervals based on bootstrap resampling. Positive values denote parameters that increase mosquito abundance, while negative values indicate parameters that reduce abundance.

Several limitations of this study should be acknowledged. First, the absence of mosquito surveillance data for the study area prevented direct comparison of model outputs with field observations, and our conclusions are therefore based on theoretical projections. We hope that this work will encourage systematic collection of both aquatic and adult mosquito data across sites to support future model validation. Second, some key parameters, including number of eggs laid per oviposition (*b*) and adult diapause mortality (*µ*_*Ad*_) lack robust fieldbased estimates; we assumed constant values within literature-informed ranges, which may not fully capture site-specific variability. Third, rainfall-dependent parameters were parameterized using functional relationships from previous studies in the absence of laboratory data, introducing additional uncertainty. Finally, we assumed that mosquito mortality rates are primarily temperature-dependent, with rainfall effects represented indirectly through development rates rather than explicitly modeled. Empirical studies quantifying rainfall impacts on mosquito mortality would strengthen future modeling efforts. Despite these limitations, the model provides valuable insights into how interacting climatic drivers shape mosquito dynamics and highlights priorities for data collection and experimental work.

## 5 Conclusion

In this study, we developed a stage-structured model of *Culex* mosquitoes that explicitly incorporates temperature, rainfall, and photoperiod to capture the seasonal and climatic regulation of population dynamics. By integrating stochastic rainfall sampling with temperature- and photoperiod-dependent functions, the model provides a mechanistic framework for exploring how interacting climate drivers shape mosquito abundance. Simulations revealed strong nonlinear responses to warming, with moderate warming intensifying the effects of rainfall differences on mosquito abundance and higher warming leading to convergence at consistently high densities. Rainfall determined whether populations were suppressed, intermediate, or amplified, while warming extended the active season and raised baseline abundance. Together, these findings emphasize that climate variability govern mosquito dynamics, with important implications for ecology and public health. The framework presented here not only clarifies the mechanisms by which environmental change influences vector populations but also offers a foundation for future extensions, including adaptive carrying capacities, refined parameterization from field and laboratory data, and explicit coupling with pathogen transmission models.

Future research should further explore the role of temperature in modifying diapause behavior, particularly under climate change. Warmer conditions may shift the timing of diapause termination or alter its intensity, with consequences for mosquito seasonality and transmission potential. Incorporating the combined effects of photoperiod and temperature on diapause into future models could therefore improve predictions of *Culex* activity and inform more targeted control strategies. In addition, our sensitivity analysis highlighted key parameters, such as number of eggs laid per oviposition and adult diapause mortality, that strongly influence population outcomes but remain poorly parameterized. Updated field and laboratory studies aimed at refining these estimates will be critical for improving model accuracy. Finally, extending this framework to consider resource-or climate-dependent carrying capacities, and coupling population dynamics with pathogen transmission models, would provide a more comprehensive understanding of how environmental change shapes vector-borne disease risk.

## Statements and Declarations

Amy Hurford acknowledges financial support from the Natural Sciences and Engineering Research Council of Canada (NSERC) through Discovery Grant RGPIN RGPIN-2023-05905.

## Data Availability Statement

The climate and weather datasets (temperature, rainfall, and photoperiod) analyzed in this study are publicly available. Daily climate data were obtained from the [Environment Canada Climate Data portal](https://climate.weather.gc.ca/climate_data/daily_data_e.html) and the [Government of Canada Climate Data portal](https://climate-change.canada.ca/climate-data/#/daily-climate-data). Monthly averaged weather history data were cross-checked against the World Weather Online archive. The R code used for data processing and analysis is openly available on GitHub at https://github.com/jbaafi/Climate-Data-Analysis. The simulation code and outputs generated during the study are available from the corresponding author upon reasonable request.

## A Appendix

### A.1 Model formulation and description

Our mathematical model for *Culex* species population dynamics incorporates temperature- and rainfall-dependent development rates, temperature-dependent oviposition and mortality rates, photoperiod-dependent diapause induction and termination, and density-dependent mortality in the larval stage. Temperature, rainfall, and photoperiod are all treated as time-dependent environmental drivers.

Let *E*(*t*), *L*(*t*), *P* (*t*), *A*(*t*), and *A*_*d*_(*t*) denote the abundances of eggs, larvae, pupae, active adults, and diapausing adults at time *t*, respectively. Assuming that adult females mate upon emergence from the pupal stage, we describe the following stage-structured system of ordinary differential equations,

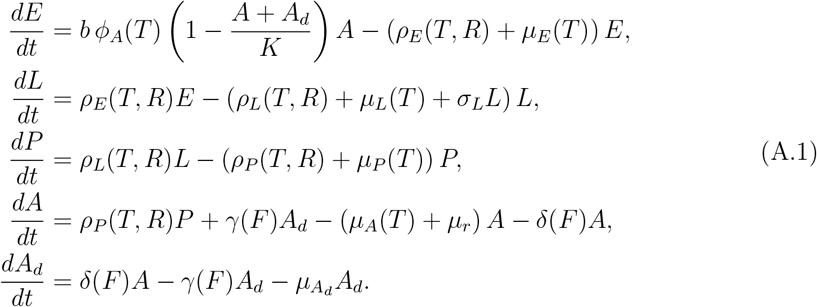

The term 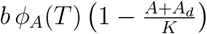 represents the density-dependent oviposition rate of active adult females, where *K* is the environmental carrying capacity of adult mosquitoes, *A* + *A*_*d*_ is the total adult population, and *ϕ*_*A*_(*T*) is the temperature-dependent fecundity function (Mordecai, Caldwell, et al., 2019; Hancock et al., 2016). Eggs hatch into larvae at a rate *ρ*_*E*_(*T, R*) that depends on both temperature and rainfall (Rueda et al., 1990; Bomblies, Duchemin, and Eltahir, 2008; Tran et al., 2013). Larvae mature into pupae at a temperature- and rainfall-dependent rate *ρ*_*L*_(*T, R*), and pupae develop into active adults at a corresponding rate *ρ*_*P*_ (*T, R*) (Rueda et al., 1990; Paaijmans et al., 2010; Bomblies, 2012). These juvenile development rates depend on both temperature and rainfall because temperature strongly influences developmental time, while rainfall governs the availability of aquatic habitats required for immature stages (Mordecai, Paaijmans, et al., 2013; Mordecai, Cohen, et al., 2017). Extreme conditions, such as very high or low temperatures, reduce survival and slow development, while excessive rainfall can prolong larval development time due to habitat flushing (Paaijmans et al., 2010). Active adults *A* emerge from pupae and can either reproduce or enter diapause depending on environmental cues (Denlinger and Armbruster, 2014; Nelms et al., 2013). Adults reproduce at a rate *b ϕ*_*A*_(*T*), but experience an additional mortality *µ*_*r*_, representing risks associated with host-seeking and blood-feeding (Brady et al., 2013; Reisen, Milby, and Meyer, 1992). Mortality of active adults is otherwise captured by *µ*_*A*_(*T*), which varies with temperature (Yang et al., 2009; Mordecai, Caldwell, et al., 2019; Brady et al., 2013). Density dependence is also expressed at the larval stage through the crowding term *σ*_*L*_*L*, which increases larval mortality as larval density rises (Abdelrazec and Gumel, 2017; Reiskind and Janairo, 2015). Diapause is modeled explicitly through the states *A* (active adults) and *A*_*d*_ (diapausing adults) (Lou et al., 2019; Frantz et al., 2024; Denlinger and Armbruster, 2014). Entry into diapause occurs at a rate *δ*(*F*), driven primarily by photoperiod *F*, while termination of diapause occurs at a rate *γ*(*F*), again determined by photoperiod (Frantz et al., 2024; Ewing et al., 2016; Nelms et al., 2013). Once in diapause, adults experience an additional mortality 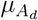, reflecting the survival costs of overwintering (Mitchell, 1981; Frantz et al., 2024). The inclusion of diapause allows the model to capture seasonal carryover of adult populations across unfavorable periods, a key feature of temperate *Culex* dynamics (Ewing et al., 2016; Frantz et al., 2024; Nelms et al., 2013).

Together, this formulation links the mosquito life cycle with environmental drivers: temperature modulates development, fecundity, and survival; rainfall regulates juvenile habitat availability and photoperiod governs diapause entry and termination (Beck-Johnson et al., 2013; Frantz et al., 2024; Parham and Michael, 2010). By explicitly embedding these processes in the system of ODEs, the model captures nonlinear responses of mosquito populations to interacting climate drivers and provides a framework for exploring ecological thresholds and climate-sensitive periods of mosquito activity.

### A.2 Simulation and analysis

All simulations and statistical analyses were performed in `R` (version 4.4.3). The system of ODEs (Appendix A.1) was solved using the `deSolve` package, with the `ode23` solver applied at a daily time step. Each climate scenario (temperature shift × rainfall regime) was run with multiple stochastic rainfall replicates to capture variability. Unless otherwise specified, analyses focused on years 3–5 of the simulations to minimize transient effects from initial conditions. For each scenario, the primary output variables (e.g., stage-specific abundances, active adult abundance, total adult abundance) were aggregated across replicates to compute summary statistics, including the mean, standard deviation, and 95% prediction intervals. Prediction intervals were estimated using the bootstrap method with 1000 resamples (Efron and Tibshirani, 1994).

Sensitivity analysis (Appendix A.4) was performed using Latin hypercube sampling (LHS) combined with partial rank correlation coefficients (PRCCs) to quantify the influence of parameters on adult abundance. PRCCs and their 95% confidence intervals were computed using the `sensitivity` package in R (Iooss et al., 2021).

Figures were produced using the `ggplot2` package (Wickham, 2016), with layouts and annotations composed using the `patchwork` package (Pedersen, 2019). Time-series plots displayed all replicate trajectories in grey to illustrate stochastic variability, with the mean trajectory overlaid in colour. Violin plots, boxplots, and scatter plots were used to summarize distributions of abundance and sensitivity results. All axes, legends, and annotations were formatted for clarity and consistency with journal style. To ensure reproducibility, all code was version-controlled and simulation scripts are available upon request.

### A.3 Data

#### A.3.1 Weather data

Daily weather observations were obtained from the St. John’s A weather station (Environment Canada) for the period 2000–2012, which provided the longest continuous record available at this site. Historical daily climate data are publicly accessible from the Environment Canada Climate Data portal^1^. Additional daily climate data were retrieved from the Government of Canada’s Climate Data portal^2^, and monthly averaged weather history data were cross-checked against the World Weather Online archive^3^.

#### Temperature

Daily mean temperature records were available for 2008–2023, excluding 2012 due to substantial missing values. These data were averaged across years to compute the mean daily seasonal cycle. A periodic cosine function of the form,

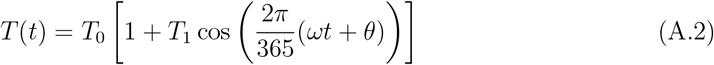

was then fitted to the averaged series, providing a smooth representation of seasonal temperature variation. The fitted curve was subsequently used to drive temperature-dependent processes in the model, including development, survival, and reproduction (See Section 2.2 and 2.3).

#### Rainfall

Daily total rainfall data exhibited high variability, precluding the use of a simple periodic function. To capture this variability while preserving observed seasonality, rainfall was treated as a stochastic input. For each simulation day, a rainfall value was randomly sampled (with replacement) from historical records for the same calendar month across the period 2008–2023. This approach preserved the monthly rainfall distribution while introducing stochastic variability, reflecting the episodic and unpredictable nature of rainfall events.

#### Photoperiod

Photoperiod data for St. John’s, NL, were obtained programmatically using the `suncalc` package in R, which calculates sunrise and sunset times for a specified geographic location given latitude and longitude. The `getSunlightTimes` function was used to compute daily sunrise and sunset times, and photoperiod length was derived as the difference between the two. Average daily photoperiod was then calculated for the period 2008–2023. To capture the seasonal pattern, a periodic cosine function was fitted to the averaged series,

The fitted photoperiod curve was used to drive diapause induction and termination processes in the model (See Section 2.2 and 2.3).

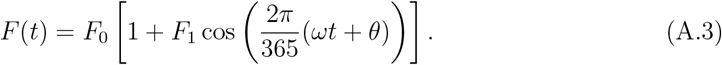

#### A.3.2 Estimating Diapause Mortality Rate of *Culex*

In the main text (Section 2.3.1), the daily mortality rate of diapausing adults 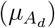 was introduced as a constant parameter, reflecting survival costs during overwintering. To estimate this rate, we relied on existing studies indicating that approximately 55% of diapausing *Culex* females survive a five-month overwintering period (≈ 167 days) under temperate conditions (Koenraadt et al., 2019; Frantz et al., 2024; Newfoundland and Labrado, 1998). Assuming constant daily mortality, survival over 167 days is modeled as,

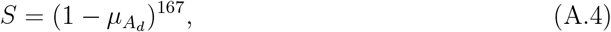

where *S* = 0.55 denotes the proportion of survivors. Solving for 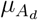 yields

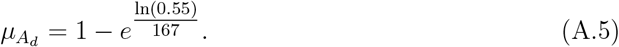

Substituting values, ln(0.55) *≈ −*0.5978 and 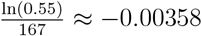, giving

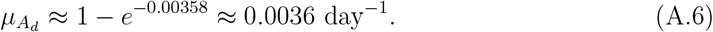

Thus, the estimated daily mortality of diapausing adults is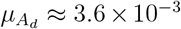, corresponding to an average survival probability of roughly 99.6% per day. This value is used in the model (Eq. A.1) to represent the additional mortality experienced by overwintering females.

### A.4 Global sensitivity analysis

To complement the simulation results, we performed a global sensitivity analysis on constant model parameters using Latin hypercube sampling (LHS) and partial rank correlation coefficients (PRCCs) with bootstrap confidence intervals (see Section A.2). The results (Figure 10) show that number of eggs laid per oviposition (*b*) and adult diapause mortality 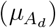 exert the strongest influence on mean adult abundance, while other parameters have weaker effects. These findings support the robustness of our conclusions and emphasize the dominant role of climate drivers (temperature, rainfall, and photoperiod) in shaping mosquito population outcomes.

### A.5 Table of Parameter Values

## Footnotes

1 Environment Canada Climate Data

2 Government of Canada: Daily Climate Data

3 World Weather Online

## Notes

https://climate.weather.gc.ca/climate_data/daily_data_e.html

https://climate-change.canada.ca/climate-data/\#/daily-climate-data

https://www.worldweatheronline.com/saint-johns-weather-averages/newfoundland-and-labrador/ca.aspx

### Competing Interest Statement

The authors have declared no competing interest.

## References

Abbasi, Ebrahim (2025). The impact of climate change on Aedes aegypti distribution and dengue fever prevalence in semi-arid regions: A case study of Tehran Province, Iran. In: Environmental Research 275, p. 121441.

Abdelrazec, Ahmed and Abba B Gumel (2017). Mathematical assessment of the role of temperature and rainfall on mosquito population dynamics. In: Journal of mathematical biology 74.6, pp. 1351–1395.

Baafi, J (2025). Flow diagram of mosquito population dynamics. Created with BioRender.com. Accessed: 2025-09-05. URL: https://biorender.com/5ir7tjm.

Baafi, Joseph and Amy Hurford (2025). Modeling the Impact of Seasonality on Mosquito Population Dynamics: Insights for Vector Control Strategies. In: Bulletin of Mathematical Biology 87.2, p. 33.

Beck-Johnson, Lindsay M et al. (2013). The effect of temperature on Anopheles mosquito population dynamics and the potential for malaria transmission. In: PLOS one 8.11, e79276.

Beck-Johnson, Lindsay M (2017). The importance of temperature fluctuations in understanding mosquito population dynamics and malaria risk. In: Royal Society open science 4.3, p. 160969.

Bhatt, Samir et al. (2013). The global distribution and burden of dengue. In: Nature 496.7446, pp. 504–507. doi: 10.1038/nature12060. URL: https://doi.org/10.1038/nature12060.

Bhowmick, Suman et al. (2025). A weather-driven mathematical model of Culex population abundance and the impact of vector control interventions. In: Ecological Informatics, p. 103163.

Cailly, Priscilla et al. (2012). A climate-driven abundance model to assess mosquito control strategies. In: Ecological Modelling 227, pp. 7–17.

Centers for Disease Control and Prevention (2024). About Eastern Equine Encephalitis. https://www.cdc.gov/eastern-equine-encephalitis/about/index.html?CDC_AAref_Val=https://www.cdc.gov/easternequineencephalitis/statistics/index.html.

Centers for Disease Control and Prevention (2025). Data and Maps for West Nile. https://www.cdc.gov/west-nile-virus/data-maps/index.html.

Ciota, Alexander T and Alexander C Keyel (2019). The role of temperature in transmission of zoonotic arboviruses. In: Viruses 11.11, p. 1013.

Ciota, Alexander T, Amy C Matacchiero, et al. (2014). The effect of temperature on life history traits of Culex mosquitoes. In: Journal of medical entomology 51.1, pp. 55–62.

Coon, Kerri L, Shivanand Hegde, and Grant L Hughes (2022). Interspecies microbiome transplantation recapitulates microbial acquisition in mosquitoes. In: Microbiome 10.1, p. 58.

Diniz, Diego Felipe Araujo et al. (2017). Diapause and quiescence: dormancy mechanisms that contribute to the geographical expansion of mosquitoes and their evolutionary success. In: Parasites & vectors 10.1, p. 310.

Ewing, David Andrew et al. (2016). Modelling the effect of temperature on the seasonal population dynamics of temperate mosquitoes. In: Journal of theoretical biology 400, pp. 65–79.

Ezanno, Pauline et al. (2015). A generic weather-driven model to predict mosquito population dynamics applied to species of Anopheles, Culex and Aedes genera of southern France. In: Preventive veterinary medicine 120.1, pp. 39–50.

Farajollahi, A. et al. (2011). Culex pipiens complex in North America: distribution, ecology, and disease vector potential. In: Journal of the American Mosquito Control Association 27.4, pp. 1–24.

Frantz, Rachel M et al. (2024). Age structured partial differential equations model for Culex mosquito abundance. In: Ecological Modelling 494, p. 110764.

Githeko, Andrew K. et al. (2000). Climate change and vector-borne diseases: a regional analysis. In: Bulletin of the World Health Organization 78.9, pp. 1136–1147.

Gubler, Duane J. (2011). Dengue, Urbanization and Globalization: The Unholy Trinity of the 21st Century. In: Tropical Medicine and Health 39.4 Suppl, pp. 3–11. doi: 10.2149/tmh.2011-S05. URL: https://doi.org/10.2149/tmh.2011-S05.

IPCC (2021). Summary for Policymakers. In: Climate Change 2021: The Physical Science Basis. Contribution of Working Group I to the Sixth Assessment Report of the Intergovernmental Panel on Climate Change. Ed. by V. Masson-Delmotte et al. Cambridge, United Kingdom and New York, NY, USA: Cambridge University Press, pp. 3–32. doi: 10.1017/9781009157896.001.

Jia, Pengfei et al. (2016). A climate-driven mechanistic population model of Aedes albopictus with diapause. In: Parasites & vectors 9.1, p. 175.

Jobling, B (1938). On two Subspecies of Culex pipiens L.(Diptera). In: Transactions of the Royal Entomological Society of London 87.10. Accessed: 2025-09-05, pp. 193–209.

Koenraadt, Constantianus JM et al. (2019). Effect of overwintering on survival and vector competence of the West Nile virus vector Culex pipiens. In: Parasites & vectors 12, pp. 1–9.

Lindsey, N.P., J.E. Staples, and M. Fischer (2018). Eastern equine encephalitis virus disease in the United States, 2003–2016. In: American Journal of Tropical Medicine and Hygiene 98.6, pp. 1629–1634.

Lutambi, Angelina Mageni et al. (2013). Mathematical modelling of mosquito dispersal in a heterogeneous environment. In: Mathematical biosciences 241.2, pp. 198–216.

Madder, DJ, GA Surgeoner, and BV Helson (1983). Induction of diapause in Culex pipiens and Culex restuans (Diptera: Culicidae) in southern Ontario. In: The Canadian Entomologist 115.8, pp. 877–883.

Mordecai, Erin A et al. (2019). Thermal biology of mosquito-borne disease. In: Ecology letters 22.10, pp. 1690–1708.

Morin, Cory W, Andrew C Comrie, and Kacey Ernst (2013). Climate and dengue transmis-sion: evidence and implications. In: Environmental health perspectives 121.11-12, pp. 1264–1272.

Newfoundland, Heritage and Labrado (1998). Winter. URL: https://www.heritage.nf.ca/articles/environment/seasonal-winter.php#:∼:text=Both%20the%20depth%20and%20duration,until%20early%20May%2C%20with%20late%2D (visited on 03/10/2025).

Okuneye, Kamaldeen, Ahmed Abdelrazec, and Abba B Gumel (2018). Mathematical analysis of a weather-driven model for the population ecology of mosquitoes. In: Mathematical Biosciences & Engineering 15.1, pp. 57–93.

Pan American Health Organization (2024). Epidemiological Update: Dengue in the Americas. https://www.paho.org/en/documents/epidemiological-update-dengue-americas-june-2024.

Pan American Health Organization (2025). PAHO warns of increased risk of dengue outbreaks due to circulation of DENV-3 in the Americas. https://www.paho.org/en/news/10-2-2025-paho-warns-increased-risk-dengue-outbreaks-due-circulation-denv-3-americas.

Parham, Paul E. and Edwin Michael (2009). Modeling the effects of weather and climate change on malaria transmission. In: Environmental Health Perspectives 118.5, pp. 620–626. doi: 10.1289/ehp.0901256.

Petersen, L.R., A.C. Brault, and R.S. Nasci (2013). West Nile Virus: Review of the Literature. In: JAMA 310.3, pp. 308–315.

Reinhold, Joanna M, Claudio R Lazzari, and Chloé Lahondère (2018). Effects of the environmental temperature on Aedes aegypti and Aedes albopictus mosquitoes: a review. In: Insects 9.4, p. 158.

Reisen, W.K. (2013). Ecology of West Nile virus in North America. In: Viruses 5.8, pp. 2079–2105.

Reiter, Paul (2001). Climate change and mosquito-borne disease. In: Environmental health perspectives 109.uppl 1, pp. 141–161.

Ryan, Sadie J et al. (2019). Global expansion and redistribution of Aedes-borne virus transmission risk with climate change. In: PLoS neglected tropical diseases 13.3, e0007213.

Soetaert, Karline, Thomas Petzoldt, and R. Woodrow Setzer (2010). Solving Differential Equations in R: Package deSolve. In: Journal of Statistical Software 33.9, pp. 1–25. doi: 10.18637/jss.v033.i09. URL: https://www.jstatsoft.org/article/view/v033i09.

Tolle, Michael A. (2009). Mosquito-borne Diseases. In: Current Problems in Pediatric and Adolescent Health Care 39.4, pp. 97–140. doi: 10.1016/j.cppeds.2009.01.001. URL: https://doi.org/10.1016/j.cppeds.2009.01.001.

Weaver, Scott C. et al. (2018). Zika, Chikungunya, and Other Emerging Vector-Borne Viral Diseases. In: Annual Review of Medicine 69, pp. 395–408. doi: 10.1146/annurev-med-050715-105122. URL: https://doi.org/10.1146/annurev-med-050715-105122.

Winokur, Olivia C et al. (2020). Impact of temperature on the extrinsic incubation period of Zika virus in Aedes aegypti. In: PLoS neglected tropical diseases 14.3, e0008047.

World Health Organization (2023). World Malaria Report 2023. Tech. rep. Accessed: 27 August 2025. Geneva, Switzerland: World Health Organization. URL: https://www.who.int/teams/global-malaria-programme/reports/world-malaria-report-2023.

World Health Organization(Aug. 2025). Dengue and severe dengue. Fact sheet. Accessed: 2025-09-04. URL: https://www.who.int/news-room/fact-sheets/detail/dengue-and-severe-dengue.

Yang, Guofa et al. (2009). A survey of indoor resting malaria vectors and their insecticide resistance status in Mali. In: Malaria Journal 8, p. 299. doi: 10.1186/1475-2875-8-299.

## Appendix References

Bomblies, Arne (2012). Modeling the role of rainfall patterns in seasonal malaria transmission. In: Climatic change 112.3, pp. 673–685.

Bomblies, Arne, Jean-Bernard Duchemin, and Elfatih AB Eltahir (2008). Hydrology of malaria: Model development and application to a Sahelian village. In: Water Resources Research 44.12.

Brady, Oliver J et al. (2013). Modelling adult Aedes aegypti and Aedes albopictus survival at different temperatures in laboratory and field settings. In: Parasites & vectors 6.1, p. 351.

Ciota, Alexander T et al. (2014). The effect of temperature on life history traits of Culex mosquitoes. In: Journal of medical entomology 51.1, pp. 55–62.

Clements, Alan Neville (1999). The biology of mosquitoes. Volume 2: sensory reception and behaviour. Accessed: 2025-09-05. Wallingford, UK: CABI Publishing. ISBN: 978-0-85199-313-3.

Denlinger, David L and Peter A Armbruster (2014). Mosquito diapause. In: Annual review of entomology 59, pp. 73–93.

Efron, Bradley and Robert J Tibshirani (1994). An introduction to the bootstrap. Chapman and Hall/CRC.

Hancock, Penelope A et al. (2016). Density-dependent population dynamics in Aedes aegypti slow the spread of wM el Wolbachia. In: Journal of Applied Ecology 53.3, pp. 785–793.

Hilker, Frank M and Frank H Westerhoff (2007). Preventing extinction and outbreaks in chaotic populations. In: The American Naturalist 170.2, pp. 232–241.

Iooss, Bertrand et al. (2021). Sensitivity: global sensitivity analysis of model outputs. In: R package version 1.0.

Lou, Yijun et al. (2019). Modelling diapause in mosquito population growth. In: Journal of mathematical biology 78.7, pp. 2259–2288.

Madder, DJ, GA Surgeoner, and BV Helson (1983a). Induction of diapause in Culex pipiens and Culex restuans (Diptera: Culicidae) in southern Ontario. In: The Canadian Entomologist 115.8, pp. 877–883.

Madder, DJ, GA Surgeoner, and BV Helson (1983b). Number of generations, egg production, and developmental time of Culex pipiens and Culex restuans (Diptera: Culicidae) in southern Ontario. In: Journal of medical entomology 20.3, pp. 275–287.

Mitchell, Carl J (1981). Diapause termination, gonoactivity, and differentiation of host-seeking behavior from blood-feeding behavior in hibernating Culex tarsalis (Diptera: Culicidae). In: Journal of medical entomology 18.5, pp. 386–394.

Mordecai, Erin A, Jamie M Caldwell, et al. (2019). Thermal biology of mosquito-borne disease. In: Ecology letters 22.10, pp. 1690–1708.

Mordecai, Erin A, Jeremy M Cohen, et al. (2017). Detecting the impact of temperature on transmission of Zika, dengue, and chikungunya using mechanistic models. In: PLoS neglected tropical diseases 11.4, e0005568.

Mordecai, Erin A, Krijn P Paaijmans, et al. (2013). Optimal temperature for malaria transmission is dramatically lower than previously predicted. In: Ecology letters 16.1, pp. 22–30.

Nelms, Brittany M et al. (2013). Overwintering biology of Culex (Diptera: Culicidae) mosquitoes in the Sacramento valley of California. In: Journal of medical entomology 50.4, pp. 773–790.

Paaijmans, Krijn P et al. (2010). Influence of climate on malaria transmission depends on daily temperature variation. In: Proceedings of the National Academy of Sciences 107.34, pp. 15135–15139.

Parham, Paul Edward and Edwin Michael (2010). Modeling the effects of weather and climate change on malaria transmission. In: Environmental health perspectives 118.5, pp. 620–626.

Pedersen, Thomas Lin (2019). Patchwork: The composer of plots. In: CRAN: Contributed Packages.

Reisen, William K, Marilyn M Milby, and Richard P Meyer (1992). Population dynamics of adult Culex mosquitoes (diptera: culicidae) along the Kern River, Kern County, California, in 1990. In: Journal of medical entomology 29.3, pp. 531–543.

Reiskind, Michael H and M Shawn Janairo (2015). Late-instar behavior of Aedes aegypti (Diptera: Culicidae) larvae in different thermal and nutritive environments. In: Journal of Medical Entomology 52.5, pp. 789–796.

Rueda, LM et al. (1990). Temperature-dependent development and survival rates of Culex quinquefasciatus and Aedes aegypti (Diptera: Culicidae). In: Journal of medical entomology 27.5, pp. 892–898.

Tran, Annelise et al. (2013). A rainfall-and temperature-driven abundance model for Aedes albopictus populations. In: International journal of environmental research and public health 10.5, pp. 1698–1719.

Turell, Michael J et al. (2005). An update on the potential of North American mosquitoes (Diptera: Culicidae) to transmit West Nile virus. In: Journal of medical entomology 42.1, pp. 57–62.

Wickham, Hadley (2016). Data analysis. In: ggplot2: elegant graphics for data analysis. Springer, pp. 189–201.

